# Combining cortical and spinal stimulation maximizes improvement of gait after spinal cord injury

**DOI:** 10.1101/2025.05.22.655593

**Authors:** Roxanne Drainville, Marco Bonizzato, Davide Burchielli, Rose Guay-Hottin, Alexandre Sheasby, Marina Martinez

## Abstract

Most spinal cord injuries (SCI) spare descending motor pathways and sublesional networks, which can be activated through motor cortex and spinal cord stimulation to mitigate locomotor deficits. However, the potential synergy between cortical and spinal stimulation as a neuroprosthetic intervention remains unknown. Here, we first investigated phase-locked electrical stimulation of the motor cortex and lumbar spinal cord at 40 Hz in a rat model of unilateral SCI. Combining cortical and lumbar stimulation around the anticipated lift synergistically enhanced leg movements. When integrated into rehabilitation training, cortical stimulation proved essential for recovery of skilled locomotion. As a further refinement, we next investigated the effects of high-frequency (330 Hz) lumbar and sacral stimulation combined with cortical stimulation. Timely integration during the swing phase showed that cortical and rostral lumbar stimulations enhance the initial and mid-swing phases, while sacral stimulation improves extension velocity in the late swing. These findings indicate that supraspinal and sublesional neuromodulation offer complementary neuroprosthetic effects in targeted SCI gait rehabilitation.

**Highlights:**

- Cortical and spinal stimulations summate motor outputs via distinct pathways.
- Each improves gait post-SCI, but combined stimulation maximizes gait improvement.
- Integrating cortico-spinal stimulation into rehabilitation promotes lasting recovery.
- EES capabilities extended using high-frequency lumbosacral protocols.

## INTRODUCTION

Spinal cord injuries (SCI) lead to persistent locomotor deficits, resulting from the disruption of pathways that connect the supraspinal centers with the spinal cord. Notably, over 70% of SCIs are incomplete ^1^, preserving some connection between the brain and the sublesional cord and offering potential for recovery through targeted rehabilitative techniques and emerging technologies.

Among the rehabilitation techniques for lower limb recovery, locomotor training has long been acknowledged for its crucial role in restoring locomotor rhythms and engaging activity-dependent spinal mechanisms ^2–4^. Moreover, increasing evidence suggests that the enhanced application of afferent input to the spinal cord, such as through electrical epidural stimulation (EES) of the lumbosacral spinal cord, may facilitate stepping activities and holds the potential to evoke reemergence of descending motor commands ^5,6^. Recent advancements include multisite EES delivery with specific locomotor phases, precisely aligning with intended movements to reproduce natural dynamics of motoneuron activation. These systems have demonstrated superiority over continuous unpatterned stimulation in restoring locomotor movements and specific muscular synergies ^7,8^.

However, the optimal EES parameters and the single impact of spinal EES over immediate movement modulation and long-term recovery of locomotor control remain poorly understood. In fact, most animal studies have combined EES with other facilitating factors, including body weight support and/or prolocomotor pharmacological agents ^9–14^, often conducted on a treadmill in the upright position (bipedal stepping) ^9–11^. These conditions are known to notably alter the excitability of spinal locomotor circuits ^15^ and enhance sensory feedback ^16^. Conversely, human large-scale adoption of EES in daily activities would be likely delivered without pharmacological enabling factors ^5,6,8^.

Both rodent ^9,10,12,14^ and human ^6,17–19^ studies consistently employ EES parameters of 25–120 Hz frequency and 0.2 ms pulse width. The most common approach is to administer EES at 40 Hz, supported by a history of studies using continuous lumbosacral EES at this frequency, indicating its optimal efficacy in facilitating leg muscle activity, enhancing kinematics, and recruiting proprioceptive afferents ^10,11,20,21^. These settings facilitate uninterrupted mono– and polysynaptic responses in the spinal locomotor circuitry ^22^ and modulate electromyographic signals from the ankle flexor and extensor muscles, promoting predictive adjustment of leg movements during locomotion ^9,10,12,14^. Here, we investigated a novel approach using high-frequency EES with the goal of increasing EES efficacy. In fact, long-duration and continuous EES can have disruptive effects by erasing proprioceptive information ^23^. Using EES with high-frequency and short duration maximizes spatiotemporal modulation of spinal circuits via their afferent inputs ^7^, which could help achieve higher efficacy in gait improvement.

Recently, neurostimulation approaches have empowered efferent drive through supraspinal stimulation to engage residual descending projections ^24–26^. Notably, a neurostimulation strategy utilizing electrical stimulation of the motor cortex has been developed to alleviate SCI deficits. When delivered in phase-coherence with the intended movement, intracortical microstimulation (ICMS) applied to the motor cortex immediately enhanced motor output and fostered recovery of cortical control of locomotion through training ^27,28^.

The coupling of spatiotemporal spinal and supraspinal approaches consequently holds potential to promote complementary locomotor control mechanisms – the former recruiting spinal locomotor circuits via sensory pathways ^23,29^ and the latter engaging residual voluntary drive ^27,30^. However, direct comparisons of the impacts of isolated cortical and spinal stimulation, as well as their combined application, on both immediate modulation and long-term locomotor recovery, are lacking.

In this study, we assessed how spatiotemporal cortical and spinal stimulations independently and synergistically modulate the kinematics of quadrupedal locomotion and comprehensively characterized the optimal parameters for maximal improvement in walking metrics following unilateral SCI in rats. In the first set of experiments, we evaluated the immediate and long-term effects of combined cortical and 40Hz spinal lumbar stimulation. We found that 40 Hz lumbar EES influenced leg kinematics in an intensity and duration-dependent manners and interacted synergistically with ICMS to alleviate locomotor deficits. Moreover, combined cortico-spinal stimulation led to superior recovery of skilled locomotion when integrated into daily locomotor training, surpassing the outcomes of rats trained with or without spinal stimulation alone. In the second phase, we expanded the capabilities of our neuroprosthesis by investigating short-duration high-frequency (330 Hz) lumbar and sacral EES and its combination with ICMS. Our findings demonstrate that integrating these three different stimulation modalities timely in the locomotor cycle collectively reinforces locomotion, facilitating a synergistic combination of flexion and extension movements.

## RESULTS

We designed a cortico-spinal neuroprosthesis (Fig. 1) integrating spatiotemporal intracortical, lumbar (L2) and sacral (S1) stimulations during locomotion, with the aim of synergistically enhancing leg flexion ^7,11,19,27,31^ and extension ^5,13,32,33^. Using a hemisection SCI model inducing paralysis in one leg and chronically affecting volitional movement control, including skilled locomotion ^34–36^, we conducted experiments before and after SCI to assess the immediate effect of ICMS and standard 40Hz L2 EES on leg kinematics, with optimized parameters for unsupported quadrupedal walking. Additionally, this cortico-spinal neuroprosthesis was employed as a neurostimulation therapy alongside locomotor training for 3 weeks to evaluate the long-term effects of combined neuroprosthetic interventions, focusing specifically on their potential to foster sustained recovery of skilled locomotion, measured through ladder task performance. Subsequently, we explored additional modalities to broaden our neuroprosthesis approach, including various L2 EES frequencies, and determined optimal parameters to maximize locomotor output when combined with S1 EES and ICMS. These evaluations were conducted without the interference of additional pharmacological or mechanical walk-aid facilitating factors, thereby establishing an intervention model capable of isolating the impact of electrical neuroprosthetic stimulation on locomotor control and recovery.

**Fig 1.**
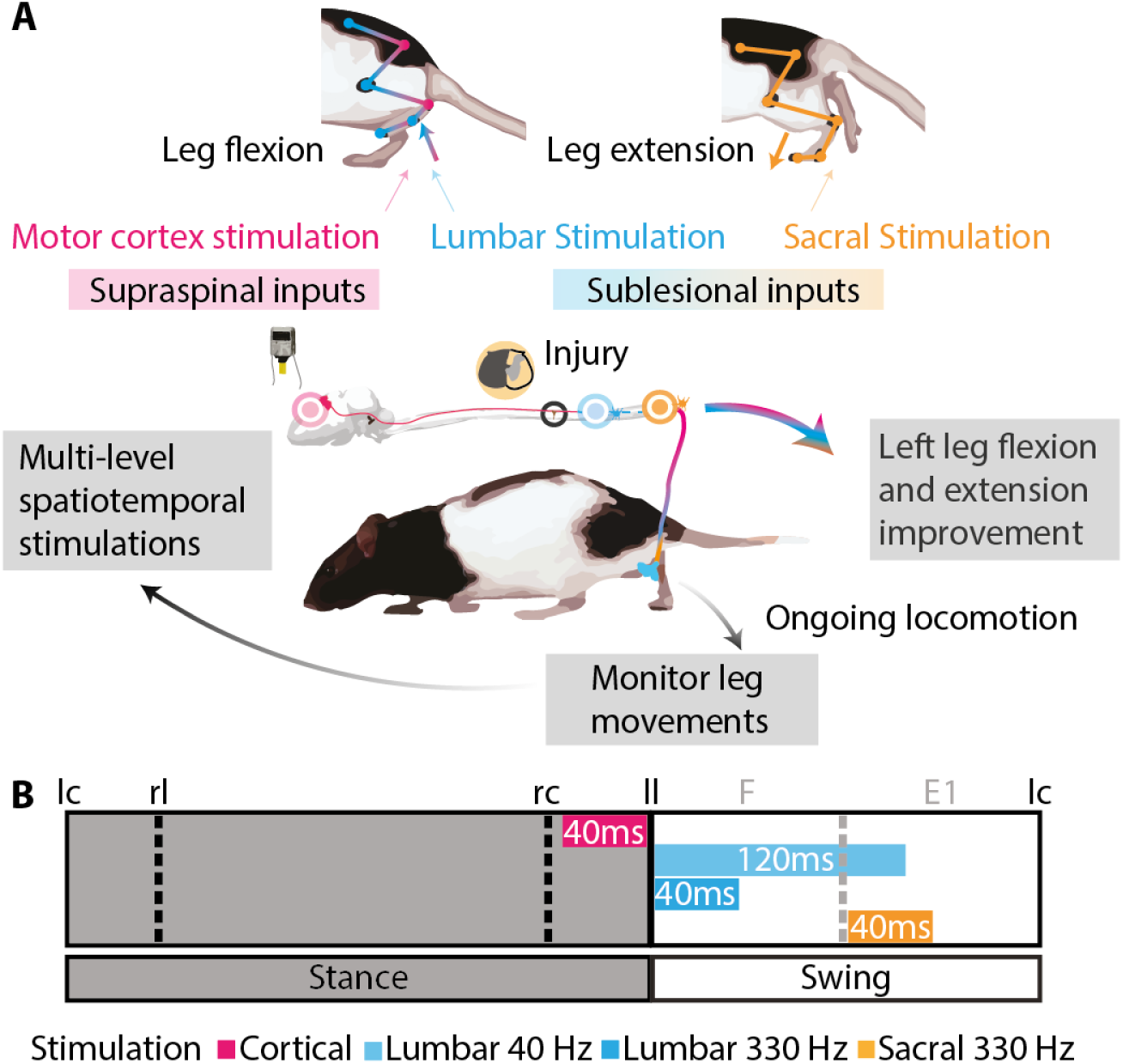
Design of the cortico-spinal neuroprosthesis. (**A**), During treadmill locomotion at 23 cm/s, electromyographic activity from hindlimb muscles was recorded and analyzed in real-time to predict the occurrence of locomotor phases with anticipation, subsequently triggering multi-level spatiotemporal stimulations. (**B**), Three sources of stimulation were delivered in synchrony with specific subphases of the gait cycle to replicate or enhance muscle activity patterns. ICMS was delivered 40 ms before the lift onset through an electrode eliciting ankle flexion within a 32-channel array. EES was then applied to the lumbar segment L2 at the lift onset. Finally, EES was applied 90 ms later at the sacral segment S1 during the second phase of the swing, corresponding to the first extension phase of the gait cycle. This sequence aimed to increase supraspinal and sublesional drive to spinal locomotor circuits, enhancing both flexion through ICMS and L2 EES and extension through S1 EES. lc: left contact, rl: right lift, rc: right contact, ll: left lift, F: flexion phase of the swing, E1: extension phase of the swing.

### Modulation of lumbar EES parameters tunes leg kinematics

We first aimed to restore flexion movements, which are significantly impaired after hemisection SCI ^34–36^. Consequently, we targeted the L2 spinal segment (Fig. 2A), known for its involvement in the generation of flexor activity ^7,11,19,31^. We applied 40Hz L2 EES in synchrony with the lift onset during treadmill walking at 23 cm/s using an electromyographic (EMG) pattern recognition algorithm capable of detecting gait phases. We considered pulse intensity (250-600 µA) (Fig. 2B) and train duration (50-250 ms) (Fig. 2C). Within this parameter range, one variable was modulated while the others were held constant. The effects of this modulation were evaluated in terms of leg movement during quadrupedal locomotion in a cohort of six rats, both in intact state and 5-10 days post-hemisection SCI. This experimental timeframe corresponds to a period during which rats exhibit a substantial lack of leg flexion, leading to dragging ^34–36^. Consequently, we identified changes in step height, flexion velocity, and the percentage of dragging as primary outcomes for this screening.

**Fig. 2:**
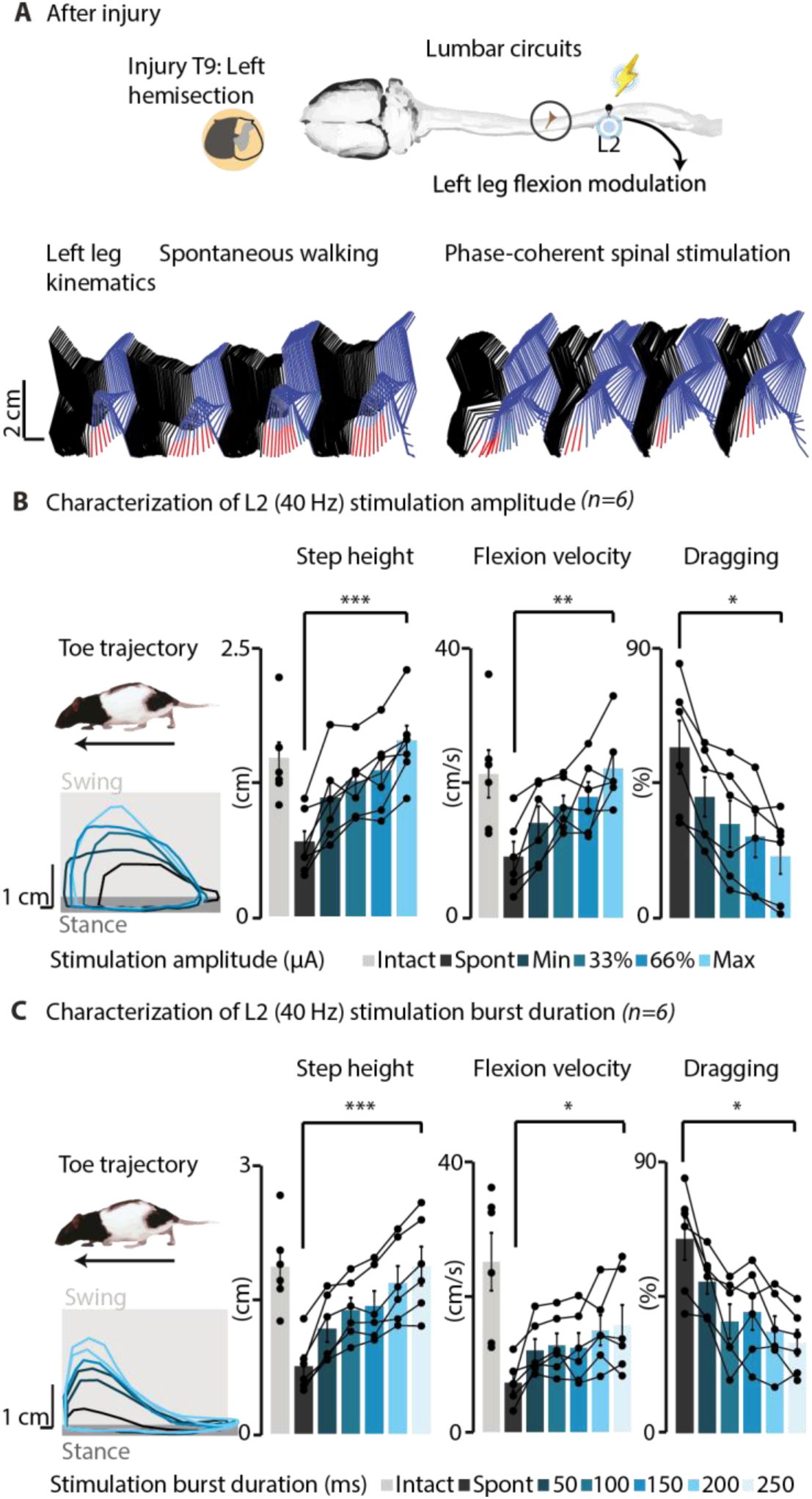
40Hz L2 EES intensity and duration modulate leg kinematics after SCI rats. (**A**), Schematic setup of EES delivery to the L2 spinal segment at the lift onset. Representative stick diagrams depict leg movements and toe trajectory without and with L2 EES in a rat (red sticks represent foot drag, blue sticks indicate the swing phase, and black sticks indicate the stance phase). (**B**), Impact of EES intensity on leg kinematics. EES intensities were randomly tested within a functional range where the minimal intensity elicited a visible muscle twitch, and the maximal intensity was set to 90% of the maximal comfortable value for each rat (250-600 µA). The burst duration was set at 120 ms. Step height modulated linearly with increased stimulation intensities (p=0.005, +160±36% of spontaneous walking, fit r^2^ min to max: 81±10%), attenuating dragging by 68±8% (p=0.028, fit r^2^: 73±24%). Flexion velocity was also enhanced with increasing intensities (p=0.007, +235±98% of spontaneous walking). (**C**), Impact of EES burst duration (50-250ms range, maximal intensity captured in **b**) on leg kinematics. An increase in EES duration resulted in a linear increase in step height (p= 0.0009, +159±27% of spontaneous walking, fit r^2^ min to max: 90±7%). Increase in flexion velocity (p=0,023, +140±53% of spontaneous walking) and decrease in dragging (p=0.028, –53±13% of spontaneous walking) were also observed. However, these parameters did not linearly correlate with EES durations (flexion velocity: fit r^2^ min to max: 49±26%; dragging %: fit r^2^ min to max: 57±18%). In all panels, individual data from n=6 rats are presented. Bar plots report mean values ± SEM. \**P*<0.05, \*\**P*<0.01, \*\*\**P*<0.001, paired, one-tailed t-test or Wilcoxon Signed Rank test.

In both the intact state (Fig. S1B) and after SCI (Fig. 2B), L2 EES intensity exhibited precise control over ipsilateral leg movements. Increasing intensity (Fig. 2B) led to a linear modulation of step height (fit r^2^ min to max: 81±10%), concurrently reducing dragging by twofold after SCI. EES duration also linearly modulated step height before (Fig. S1C) and after SCI (fit r^2^ min to max: 90±7%), increasing flexion velocity and decreasing dragging (Fig. 2C). However, further increments in EES duration beyond 100-150 ms yielded minimal gains in dragging reduction. Consequently, EES duration was consistently maintained at 120 ms in subsequent experiments using a 40 Hz frequency. This duration was chosen to effectively cover the swing duration within the step cycle without interfering with the stance phase.

### Combined cortico-spinal stimulation synergistically maximizes locomotor output

Since both L2 EES (Fig. 2) and ICMS ^27^ individually modulate leg movements in intact and SCI rats when applied in phase-coherence with the lift onset and 40 ms prior to lift onset, respectively, we investigated the potential synergistic enhancement of leg movement modulation by combining L2 EES and ICMS. Short-train ICMS (40 ms, 330 Hz) was delivered through an electrode selected from the 32-channel array implanted in the right motor cortex to produce a strong ankle flexion with the lowest threshold, since this setting minimizes charge injection and maximizes reduction of dragging ^27^. Subsequently, 40 ms later, L2 EES was applied at the lift onset (120 ms, 40Hz). This sequence allows both sources of stimulation to produce overlapping peak EMG volleys in the ankle flexor (Fig. S2A). The maximal comfortable intensity for each stimulation modality was used (Fig. S7A). The impact of L2 EES, ICMS, and their combination on leg kinematics was evaluated both before and 5-9 days after SCI in a cohort of six rats.

In both intact (Fig. S3A) and SCI (Fig. 3A) rats, the combined cortical and spinal stimulation resulted in the maximal increase in foot trajectory compared to spontaneous walking. In the intact state, the combination maximally elevated step height (Fig. S3B). Following SCI, the combined stimulation significantly improved leg kinematics compared to each individual stimulation strategy (Fig. 3B and Movie S1), resulting in increased step height and flexion velocity. Regarding dragging reduction, the cortical stimulation component proved essential for maximal alleviation of dragging deficits. These results uncovered synergies between neurostimulation strategies targeting afferent fibers projecting to lumbar spinal cord and descending supraspinal inputs orchestrated by the motor cortex and projecting to the lumbar spinal cord.

**Fig. 3:**
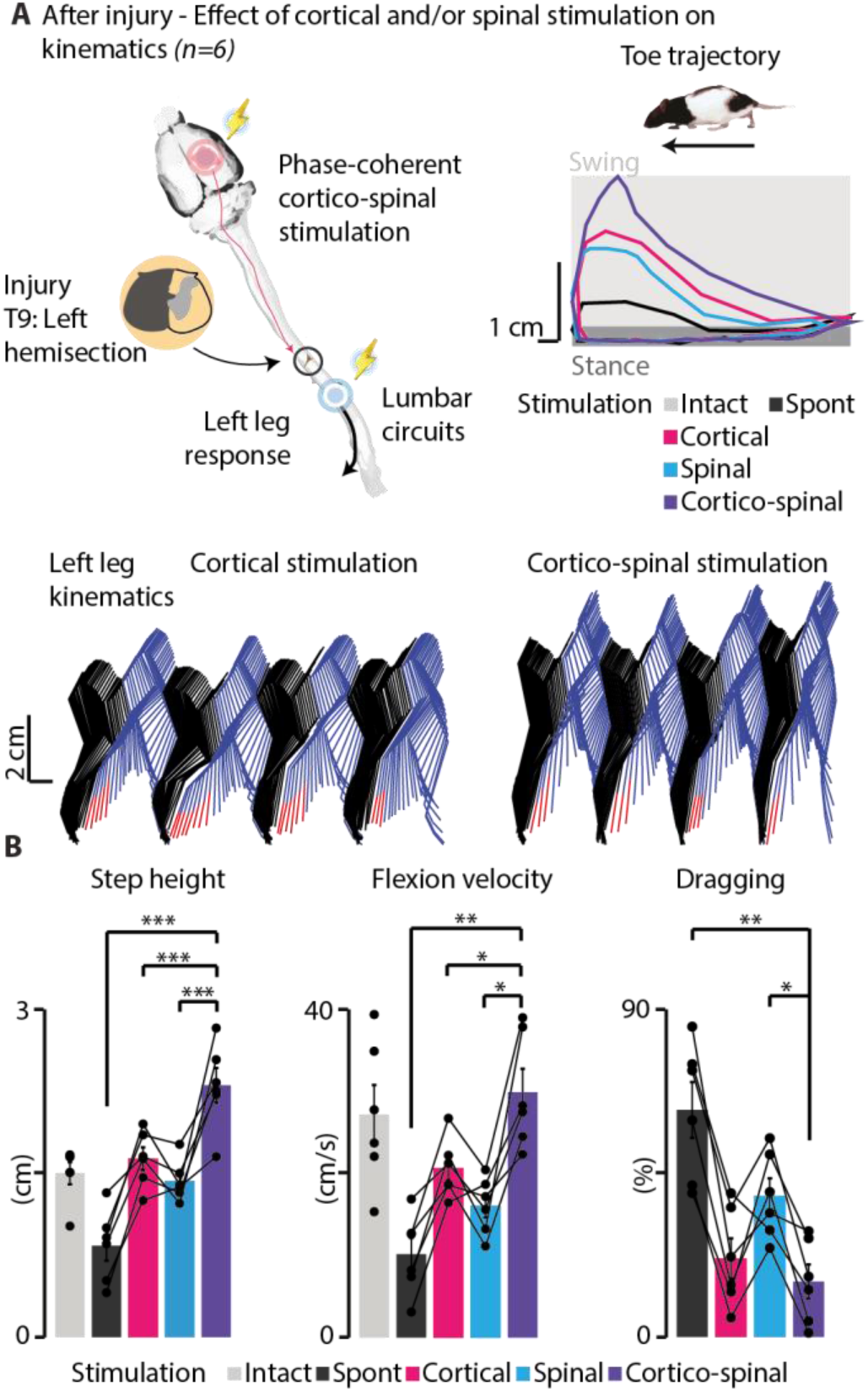
Combined cortico-spinal stimulation maximizes restoration of leg movements after SCI. (**A**), Schematic setup of cortico-spinal stimulation delivery. Representative toe trajectory and stick diagrams (red sticks represent foot drag, blue sticks indicate the swing phase, and black sticks indicate the stance phase) without and with cortical, spinal, and cortico-spinal stimulation. (**B**), Impact of individual and combined cortico-spinal stimulation on leg kinematics after SCI. Bar plots report mean values ± SEM (n=6 rats) of step height, flexion velocity and percentage of dragging for each condition of stimulation. Cortico-spinal stimulation significantly increased step height by 214±53% (p= 0.0005) and flexion velocity by 294±115% (p=0.004), concurrently reducing dragging by 75±10% (p=0.002) compared to spontaneous walking. Cortico-spinal stimulation significantly outperformed independent cortical stimulation in enhancing step height (p=0.001, +41.21±5.06%) and flexion velocity (p=0.007, +44.88±9%), but not in reducing dragging (p=0.55, ns). The combined stimulation also surpassed spinal stimulation alone in step height (p=0.007, +64±15%), flexion velocity (p=0.006, +92±23%), and reduction of dragging (p=0.008, –62±12% of spinal stimulation). \**P*<0.05, \*\**P*<0.01, \*\*\**P*<0.001, paired, one-tailed t-test or Wilcoxon Signed Rank test. The statistical significance threshold was adjusted for multiple comparisons.

### Cortico-spinal stimulation combined with rehabilitation fosters skilled locomotor recovery

Building on the superior effectiveness of cortico-spinal stimulation in improving leg motor deficits after SCI (Fig. 3) and earlier investigation demonstrating the positive influence of phase-coherent ICMS on long-term recovery of leg motor control during locomotor training ^27^, we explored the potential of maximizing locomotor recovery through combined cortico-spinal stimulation.

A total of 18 rats were divided into three groups, with 6 rats in each group. Over weeks 1 to 4 post-SCI, all rats underwent daily treadmill training (30 minutes/day) at a speed of 23 cm/s without stimulation (treadmill training group), with spinal stimulation (spinal stimulation group), or with combined cortico-spinal stimulation (cortico-spinal stimulation group) (Fig. 4A).

**Fig. 4:**
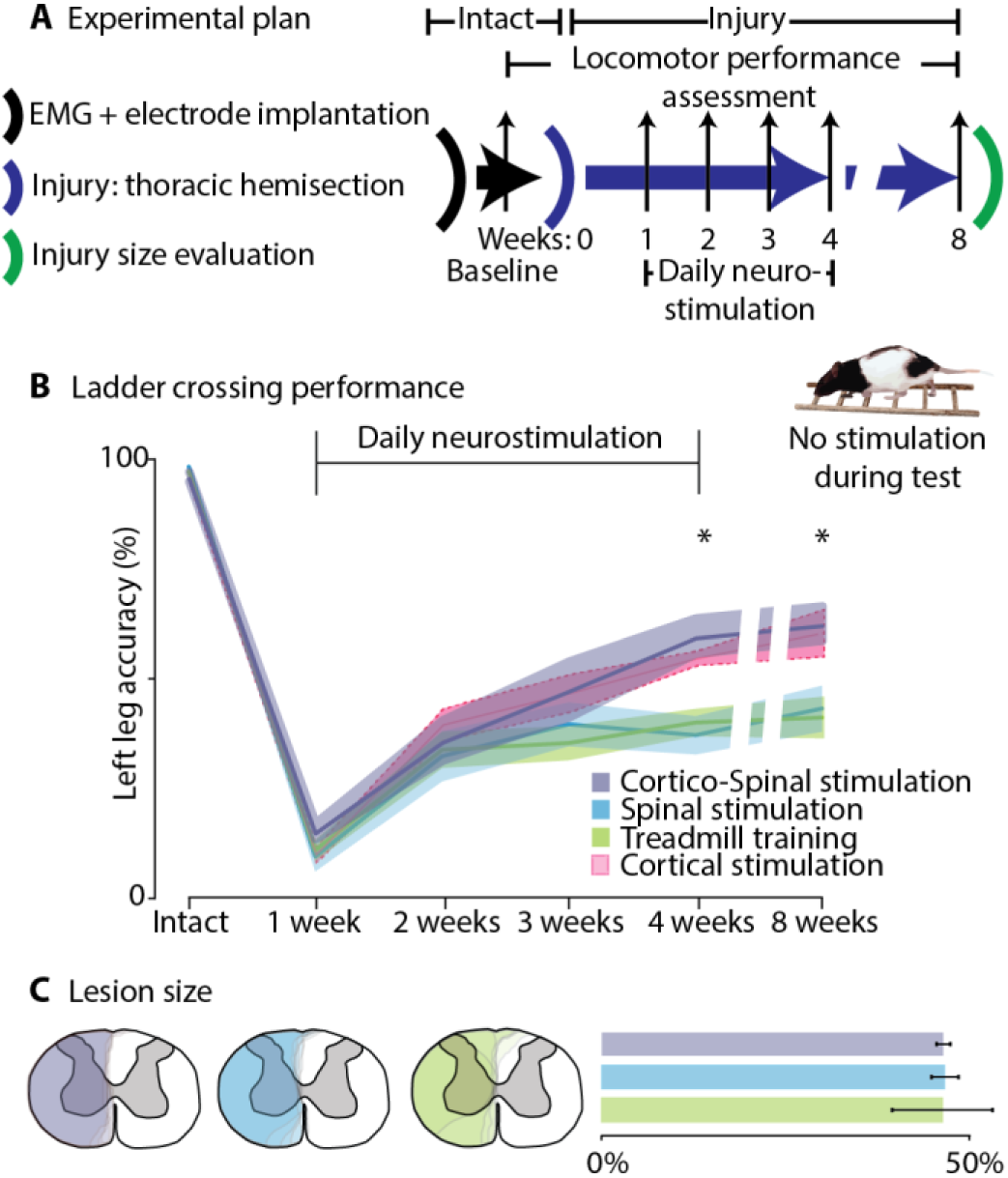
Cortico-spinal stimulation fosters recovery of skilled locomotion after SCI. (**A**), Protocol timeline. Evaluations were conducted at the intact state, followed by weekly assessments from week 1 to 4, and then again 4 weeks after therapy discontinuation. The training involved 30 min. of daily treadmill locomotion, with or without phase-coherent ICMS (40 ms, 330 Hz) and/or spinal stimulation (120 ms, 40 Hz) during weeks 1-4 after SCI. (**B**), Plots depict mean values ± SEM of ladder crossing performance. Data from rats subjected to cortical stimulation in our previous study ^27^ were re-blinded and incorporated in the plot for direct comparison, but were excluded from statistical analyses. The cortico-spinal group outperformed both the spinal stimulation and treadmill training groups at week 4 (cortico-spinal vs spinal, p=0.010; cortico-spinal vs treadmill training, p=0.03) and week 8 (cortico-spinal vs spinal, p=0.039; cortico-spinal vs treadmill training, p=0.026). *P<0.05, one-way ANOVA supplemented with Bonferroni post-hoc test. (**C**), Percentage of spared spinal tissue between groups (*P*≥0.05, one-way ANOVA).

Despite none of the stimulation strategies significantly affecting treadmill performance (Fig. S4A) and global locomotor abilities (Fig. S4B) in comparison to treadmill training alone, the recovery of skilled locomotor control, dependent on the restoration of cortical control, was facilitated by cortico-spinal stimulation. At the end of the training period (week 4) and four weeks after therapy discontinuation (week 8), rats who received cortico-spinal stimulation demonstrated superior performance compared to the spinal stimulation and treadmill training groups during ladder crossing, an untrained task conducted without any stimulation (Fig. 4B). In contrast, the performance of the treadmill training and spinal stimulation groups remained consistently comparable throughout the experiments, suggesting a limited additional benefit from spinal stimulation beyond treadmill training for cortical control recovery after unilateral SCI. The performance of the cortico-spinal stimulation group closely resembled that of the cortical stimulation group from our previous study ^27^, confirming the crucial role of cortical stimulation in the recovery of skilled locomotion. Histological analysis revealed no differences in lesion size across groups (Fig. 4C).

### High-frequency L2 EES expands stimulation capabilities

A limitation of long-duration EES is the increased risk of collisions between antidromic and natural action potentials. This may disrupt proprioceptive information ^23^, explaining the limited efficacy of continuous EES in facilitating walking after SCI in humans. In pursuit of enhanced efficacy, additional strategies were explored, including the application of short-duration, high-frequency EES. As suggested by computational models ^23,37^, such strategy may minimize proprioceptive information cancellation and expand EES efficacy. To emphasize results, we initially set L2 EES duration at 120 ms and investigated a wide frequency range (20-80 Hz) for movement modulation in intact (Fig. S5A) and SCI (Fig. S5B) rats. Notably, increasing EES frequency after SCI resulted in a linear modulation of kinematic features, such as step height (fit r^2^ min to max: 92±4%), with the highest modulation achieved at 80 Hz Fig. S5B).

We thus introduced a novel alternative approach to temporally-patterned EES, designed to mimic and match the stimulation delivery used in ICMS that is integrated with voluntary behavior ^27^. We reduced burst duration to 40 ms and delivered high-frequency L2 EES at 330 Hz. This delivery modality resulted in maximal increase in step height and flexion velocity, along with maximal decrease in dragging following SCI (Fig. S5B).

Building on these results, we investigated the timing and intensity of 330 Hz EES delivery. The most significant changes in leg kinematics in both intact (Fig. S6A) and SCI (Fig. 5A) rats were observed with EES delivered in phase-coherence with the swing. Based on the observed curves, we opted to administer EES at the lift onset and subsequently explored the impact of stimulation intensity. Our analysis revealed that kinematics exhibited a linear modulation in response to varying EES intensities, both in intact (Fig. S6B) and SCI rats (Fig. 5B).

**Fig. 5:**
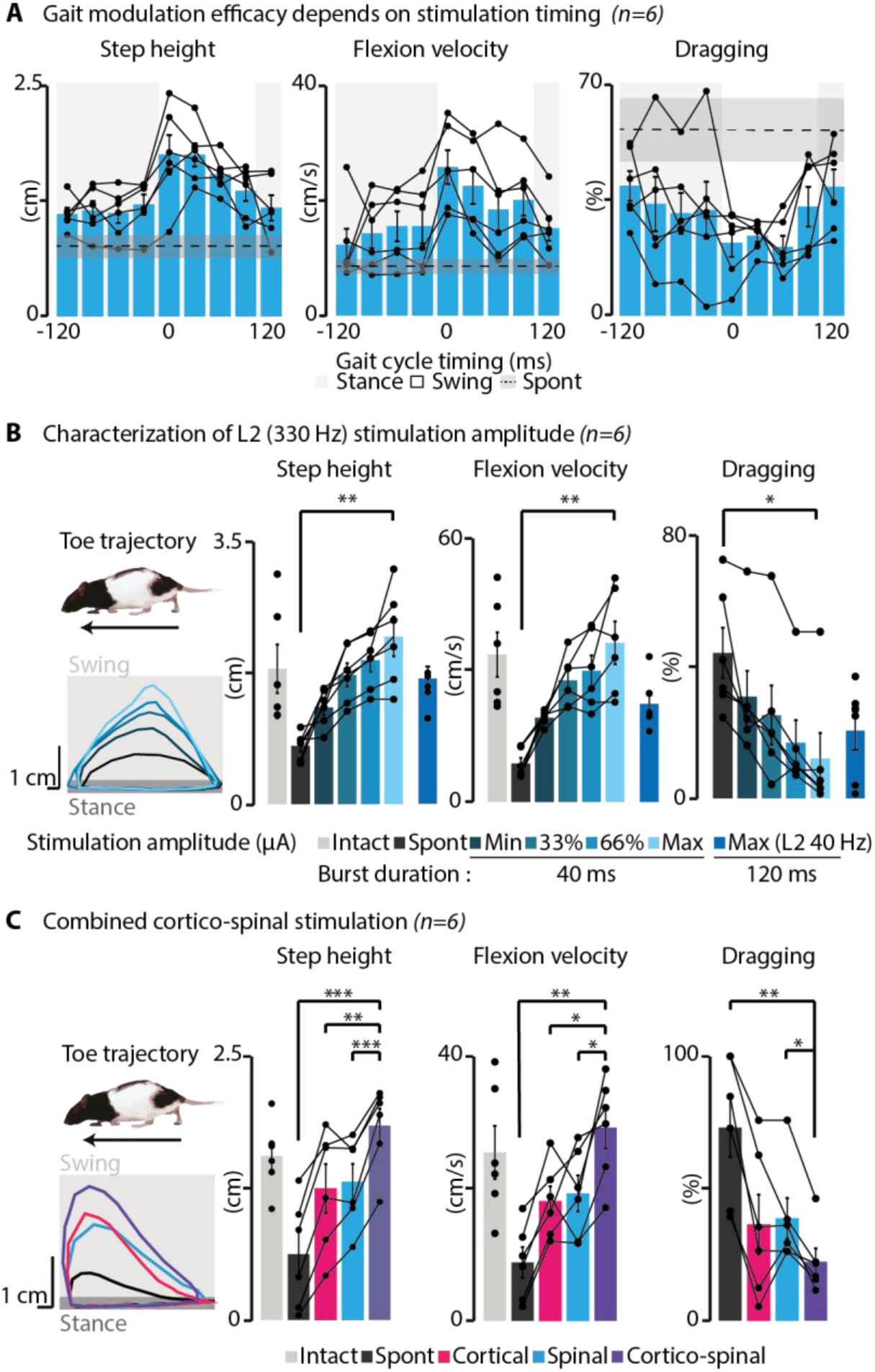
Investigating cortico-spinal stimulation capabilities using high-frequency L2 EES on movement modulation after SCI. (**A**), Impact of high-frequency L2 EES (40 ms, 330Hz) timing on leg kinematics. Step height and flexion velocity are depicted for stimulation timings ranging from 120ms before to 120 ms after the lift, with 0 representing the lift onset. The most significant increases in step height (p=0.002) and flexion velocity (p=0.002), along with a decrease in dragging (p=0.03), were obtained when EES was delivered during the swing compared to the stance phase. (**B**), Incremental increases in EES intensity (250 to 600 μA) proportionally modulated toe trajectory, leading to a linear increase in both step height (p=0.004, +225±63% of spontaneous, fit r^2^ min to max: 92±6%) and flexion velocity (p=0.004, +383±110% of spontaneous walking, fit r^2^ min to max: 73±20%), accompanied by a linear decrease in dragging (p=0.028, –81±8% of spontaneous walking, fit r^2^ min to max: 83±15%). (**C**), Combined cortical (40 ms, 330 Hz) and high-frequency EES delivered at the lift onset maximized toe trajectory, significantly increasing step height by +736±386% (p=0.0002) and flexion velocity by +443±191 (p=0.0033) compared to spontaneous walking. Compared to independent cortical and spinal stimulation, cortico-spinal stimulation increased step height by +71±25% (p=0.003) and 46±9% (p=0.0008) and improved flexion velocity by 73±28% (p=0.04) and 59±16% (p=0.02), respectively. Cortico-spinal stimulation also decreased dragging by 42±7% (p=0.011) compared to spinal stimulation. Bar plots report mean values ± SEM (n=6 rats). \**P*<0.05, \*\**P*<0.01, \*\*\**P*<0.001, paired, two-tailed t-test or Wilcoxon signed-rank test (a) and one-tailed t-test (b-c). The statistical significance threshold was adjusted for multiple comparisons. Spont: spontaneous (no stimulation).

Subsequently, we integrated this potent high-frequency EES with ICMS. L2 EES was delivered 40 ms after ICMS, aligned with the lift onset. This timing allowed both sources of stimulation to generate overlapping peak EMG volleys in the ankle flexor muscle (Fig. S2B). In both intact (Fig. S6C) and SCI (Fig. 5C and Movie S2) rats, cortico-spinal stimulation was the most effective strategy for increasing step height compared to individual EES or ICMS. Notably, in SCI rats, combined cortico-spinal stimulation outperformed spinal stimulation, increasing flexion velocity by 59±16% and reducing dragging by 42±7% compared to spinal EES alone (Fig. 5C).

Interestingly, the maximal intensities used for L2 EES at 330 Hz, whether alone or combined with ICMS, were significantly reduced compared to EES at 40 Hz in both intact and SCI rats (Fig. S7B). Additionally, the intensities were significantly lower under combined cortical and spinal stimulation at 330 Hz compared to individual L2 EES at 330 Hz or ICMS (Fig. S7C).

For an in-depth analysis of the effects of combined cortical and spinal stimulation on locomotion, we performed a multivariate analysis of kinematics parameters extracted from leg trajectories. When isolating the 2-dimensional space where the vectors indicating cortical and spinal stimulation effects lie, the combined cortico-spinal stimulation assumed an intermediate angular position, lying between the two individual stimulation vectors, both for intact (Fig. S8A) and SCI (Fig. 6A) rats, suggesting that combined stimulation benefits from the common effects of cortical and spinal gait modulation, while holding an intermediate characteristic of their unique effects. In terms of effect magnitude, the combined stimulation surpassed each of the two individual stimulations. In SCI rats, the combined stimulation brought the multivariate kinematic change closest to the projection of gait displayed by intact rats. This multivariate analysis further dissected the individual contributions of cortical and spinal stimulation effects. Specifically, EES at 330 Hz proved particularly effective in modulating early swing upward flexion, attributed to a robust early ankle closure induced by the evoked spinal reflex, leading to an increase in the elevation angle of the foot. On the other hand, cortical stimulation demonstrated effectiveness in mid-swing foot lift, promoting synergistic leg flexion, evident in a pronounced knee lift (Fig. 6B). While both types of stimulation induced distinct kinematic effects throughout the leg and successfully elevated the end effector (foot tip), only cortical stimulation had the effect to reliably trigger the swing phase (Fig. 6C, Fig. S8B). That is, cortical stimulation caused an anticipation of foot lift, while spinal stimulation did not.

**Fig. 6:**
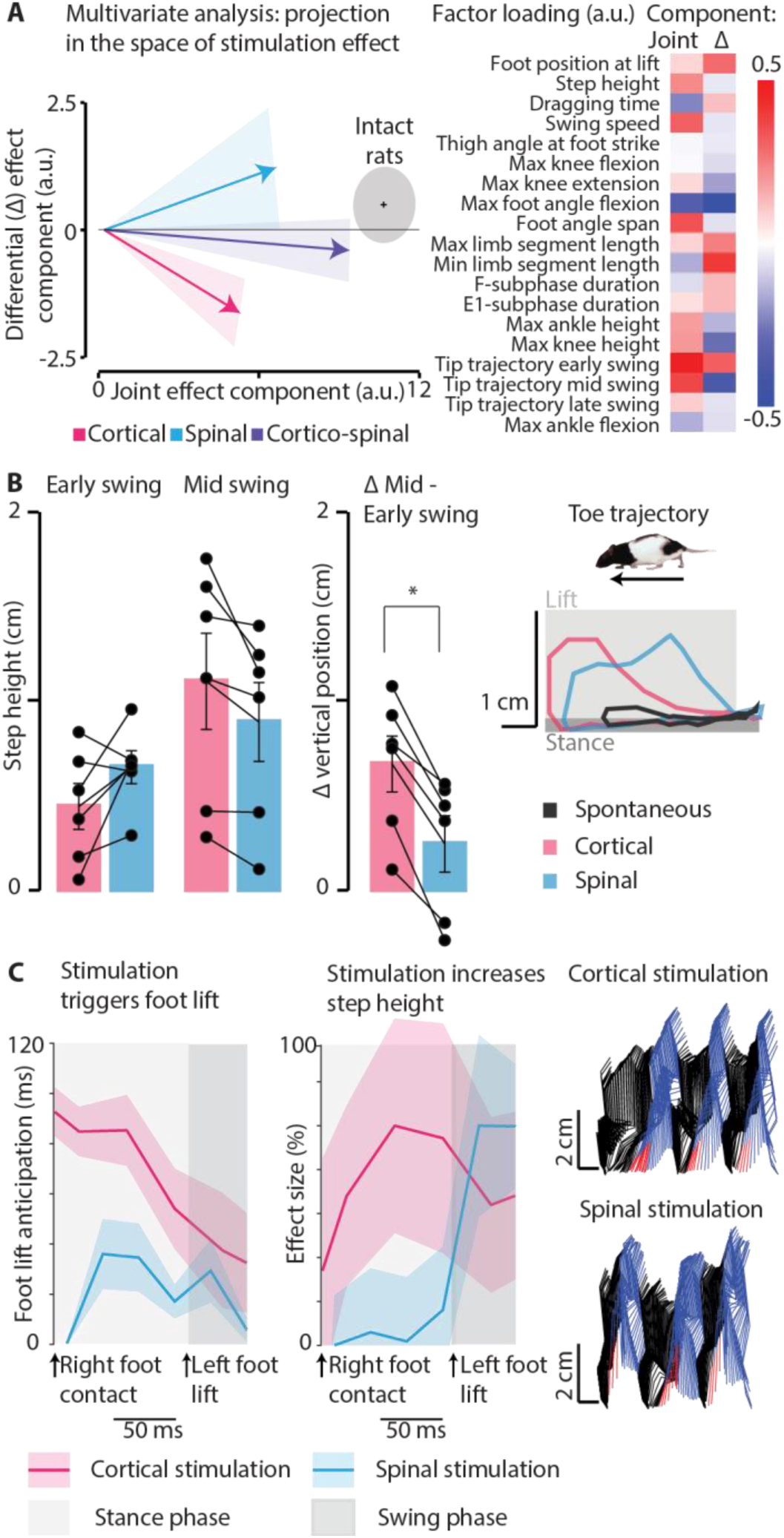
Distinct effects of cortical and spinal stimulation on gait dynamics. (**A**), Multivariate kinematic analysis qualitatively portrays enhanced gait effects of combined cortical and spinal stimulation in SCI rats. The combined cortico-spinal stimulation occupies an intermediate angular position between individual vectors, showing a greater effect magnitude and aligning SCI rats’ gait closer to intact rats (gray circle). Specific contributions include EES at 330 Hz influencing early swing upward flexion and cortical stimulation promoting mid-swing foot lift and synergistic leg flexion. (**B**), Significant differences in vertical position difference (Δ) between mid– and early swing phases were observed for trajectories shaped by cortical stimulation vs. L2 EES (330 Hz) (p=0.0013), indicating that L2 EES primarily affects early swing, while cortical stimulation exerts a stronger influence on mid-swing. (**C**), While both L2 EES and cortical stimulation increase step height, only ICMS reliably initiates the swing phase. Representative toe trajectory and stick diagrams (red sticks represent foot drag, blue sticks indicate the swing phase, and black sticks indicate the stance phase) with cortical or spinal stimulation. Error bars and shaded areas in graphs represent inter-individual variability (mean±SEM). *P<0.05, paired, two-tailed t-test.

### Integrating sacral stimulation enhances leg extension

Expanding on our findings, which demonstrate enhanced flexion movements with both cortical and high-frequency L2 EES delivered at the lift onset, we aimed to further refine the combined cortico-spinal approach. Specifically, we explored whether additional control modalities, such as sacral EES, could be integrated in the cortico-spinal neuroprosthetic platform. Our goal was to enhance extension movements, which are often impaired after SCI. To this end, we integrated sacral stimulation into our neuroprosthesis, drawing on existing research that highlights the recruitment of extensor muscle synergies through stimulation of sacral spinal segment 1 (S1) ^7,38^. ICMS was delivered first, followed 40 ms later by high-frequency L2 EES. Subsequently, 90 ms later, S1 EES was synchronized with the beginning of the extension phase of the swing, which refers to the time interval during the swing when the leg extends from maximum lift to contact with the ground (Fig. 7A). Since the latter phase is very rapid in rats, we opted for delivering a short burst (40ms) of high-frequency (330Hz) sacral stimulation. Thus, temporal patterns of cortical, L2 and S1 stimulation, except the onset delay, were identical for this experiment. Increasing S1 EES intensity modulated extension velocity by 46±6% without affecting step height (Fig. 7B).

**Fig. 7:**
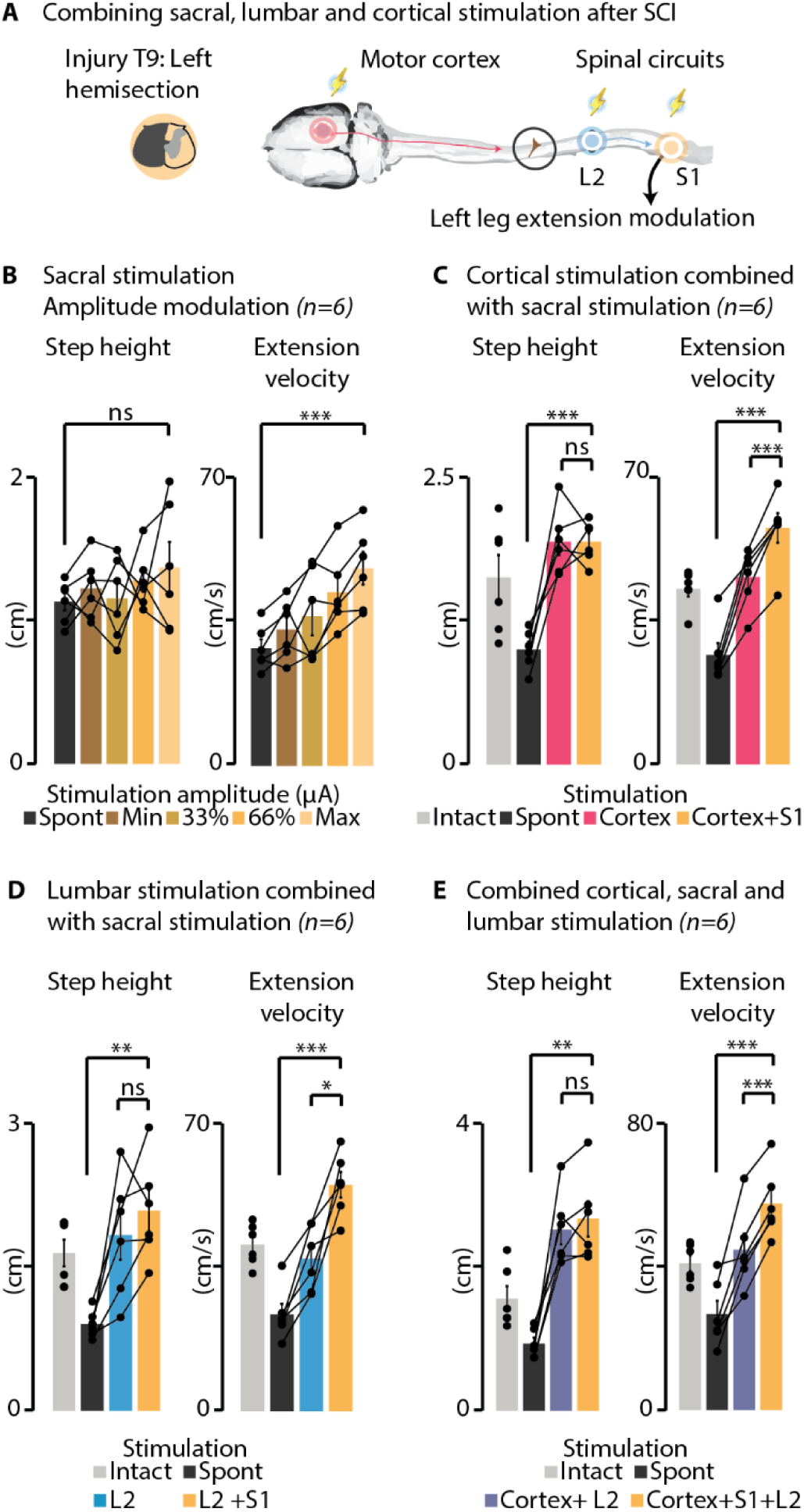
Adding sacral stimulation expands neurostimulation capabilities. (**A**), Schematic representation of the stimulation targets. Right motor cortex stimulation (40 ms, 330 Hz) was delivered first, succeeded by L2 EES (40 ms, 330 Hz) 40 ms later, and then followed by S1 EES (40 ms, 330 Hz) 90 ms after L2 EES. (**B**), Sacral stimulation modulated extension movement, resulting in a 46±6% increase in extension velocity compared to spontaneous walking (p=0.0002). When combined with cortical (**C**), L2 (**D**), or combined cortico-lumbar stimulation (**E**), S1 EES systemically increased extension velocity (124±19%, p=0.0006 compared to cortical stimulation; 141±17%, p=0.007 compared to spinal stimulation; 131±25%, p=0.00002) without impacting step height. Bar plots report mean values ± SEM (n=6 rats). \**P*<0.05, \*\**P*<0.01, \*\*\**P*<0.001, paired, one-tailed t-test. The statistical significance threshold was adjusted for multiple comparisons. Spont: spontaneous (no stimulation).

Qualitative analysis of EMG responses to isolated and combined stimulation confirmed recruitment of flexor and extensor hindlimb networks by cortical and spinal stimulation. Sacral stimulation, whereas predominantly resulting in limb extension from a kinematic perspective, are well known to have low selectivity in muscular recruitment ^39^ and this was confirmed by EMG analysis, causing flexor and extensor co-activation vastly exceeding those of L2 and cortical stimuli (Fig. S9).

Kinematically, the incorporation of S1 EES delivered during the extension sub-phase of the swing into any stimulation protocol (whether cortical, L2 or cortico-lumbar) enhanced extension velocity without further increasing the step height (Fig. 7C-E, Movie S2).

In summary, the three stimulation modalities integrate into the locomotor cycle in a phase-dependent manner, collectively reinforcing locomotion and facilitating a synergistic combination of flexion and extension movements.

## DISCUSSION

We found that cortical and lumbar spinal stimulation synergistically enhanced leg kinematics and reduced dragging. Additionally, the inclusion of sacral stimulation notably improved extension. Integrating cortico-spinal stimulation into treadmill training for 3 weeks yielded long-term recovery of voluntary motor control. We discuss the mechanisms underlying the synergetic impact of these combined neurostimulation strategies on enhancing walking and reinstating voluntary leg movements and consider the potential benefits and limitations for clinical applications.

### Integration of cortical and lumbar stimulation improves locomotion

Our findings demonstrate that both spatiotemporal ICMS and 40Hz lumbar EES immediately improve walking following SCI, with maximal alleviation of locomotor deficits observed when combining the two strategies. Previous studies indicate that spinal stepping in rodents is consistently enhanced by applying 40 Hz EES ^9,12,14^, suggesting that the stimulation frequency corresponds to an interstimulation pulse interval that enables both mono– and polysynaptic responses to occur without interruption by subsequent stimuli, possibly mirroring features of the rat’s spinal locomotor circuitry ^22^. However, when used without enabling factors, such as pharmacological stimulation or body weight support, EES induces step-like movements marked by inadequate hindlimb joint excursions, lack of coordination, and inconsistent stepping ^11,13,40,41^. Here, we present evidence of the singular efficacy of lumbar EES in manipulating foot trajectory in an intensity– and duration-dependent manner. When synchronized with the previously shown effective ICMS approach ^27,30^, it enhanced step height and flexion velocity to a greater extent. This synergistic effect likely involves additive mechanisms, where each input can reach its maximum intensity without saturating the available pool of spinal motoneurons. This aligns with studies combining supraspinal and spinal stimulation in monkeys and humans, indicating enhanced responsiveness of spinal circuits to supraspinal inputs ^42–45^. Theoretically, ICMS could be used to increase the effectiveness of EES in activating spinal locomotor circuits, potentially compensating for deficiencies of descending locomotor command pathways following SCI. Consequently, we hypothesized that daily neuroprosthetic training incorporating both forms of stimulation could optimize recovery following SCI by maximizing motor output during movement rehabilitation training or through Hebbian plasticity mechanisms. Our approach involved delivering ICMS at 330 Hz for 40 ms and EES at 40 Hz for 120 ms at lift onset, ensuring continuous EES coverage throughout the swing phase while ICMS synchronized with EES effects. Supraspinal inputs converged during the same motor phase with spinal stimulation afferent volleys in the lumbar spinal cord, but the persistent activation required to produce movement cannot be compared with existing validated paired associative stimulation protocols, characterized by brief stimuli ^46^. Our results demonstrate that cortico-spinal neuroprosthetic training was equally effective as ICMS alone in restoring voluntary motor control assessed during ladder crossing, underscoring the decisive role of cortical volleys in the restoration of volitional walking ^27,34^. In contrast to previous human ^5,6,8^ and animal ^7,10^ studies emphasizing the beneficial effects of EES on movement control recovery, we found no significant difference in ladder task performance between rats that received treadmill training alone versus those that received treadmill training combined with EES. This could be due to the nature of treadmill training, which primarily engages subcortically controlled and modulated spinal circuits, regardless of the presence of spinal stimulation ^47^. In humans, EES integrated into tasks involving volitional motor commands, such as overground walking, is pivotal for recovery ^5,6,48–50^. Priming of lumbosacral circuits through intensive neurorehabilitation paradigms has been crucial for successful strategies employing EES in both humans ^5,6,50^ and rodents ^7,10,41^. Additionally, most animal studies have employed EES as an adjunctive approach to bolster the impacts of motor training. In most cases, hindlimb stepping in rats is facilitated with EES in combination with pharmacology during non-natural bipedal movements ^10,11,51^.

Enduring benefits have been associated with an increase in cortico-spinal excitability, which is potentiated when ascending and descending volleys converge on task-relevant spinal circuits ^46,52–57^. While our cortical and spinal stimulations coincide temporally, associative plasticity hinges on the precise timing alignment between two stimulation sources at the spinal level ^43,46,58,59^, necessitating further investigation.

Therefore, interventions combining cortical and spinal approaches should aim to replicate the natural interaction between spinal stimulation and descending signals. By encouraging subjects’ voluntary efforts during these interventions, the most positive outcomes can be achieved, leading to long-term cortical and spinal cord plasticity ^5,6,50^.

### Expanding stimulation capabilities fine-tunes control of walking

Long-duration EES poses a challenge due to its potential disruption of proprioceptive feedback, which may compromise its efficacy in humans ^23^. When applied in a phase-specific manner, spinal stimulation can be pushed out of the 40-60 Hz boundaries typical for rat studies, towards higher frequencies. We explored short-duration, high-frequency spatiotemporal EES protocols ^23,37^. We found that 40 ms lumbar EES administered at 330 Hz maximally increased step height in a time– and intensity-dependent manner, with maximal effect observed at the swing onset, similar to ICMS ^27^. The reduced intensities used with high-frequency EES may have enhanced the spatial selectivity of the stimulation, as the size of the electrical field is directly related to the current intensity. However, we identified a distinct kinematic effect: ICMS effectively triggered the swing phase and had a more pronounced impact during mid-swing, resulting in a noticeable knee lift, whereas high-frequency EES primarily influenced the early swing phase, eliciting a rapid ankle closure. Cortical and spinal stimulation were then optimized to maximize lift and minimize dragging. When combined, lumbar EES and ICMS seamlessly integrated into the locomotor cycle, synergistically enhancing leg flexion throughout the swing phase. This resulted in an enhanced swing movement with amplified kinematics that exceed those achievable with either stimulation modality alone, suggesting a coordinated interplay between cortical and sensory inputs onto common spinal circuits. The increased efficacy of combined cortico-spinal stimulation in enhancing step height and flexion velocity after SCI, compared to intact animals, aligns with our previous findings suggesting that stimulation efficacy is gated by the spinal state. After hemisection SCI, alterations in the dynamics of central pattern generators result in heightened excitability within the flexor oscillator ^34,60^, potentially explaining the enhanced recruitment of flexor circuits by ICMS and lumbar EES through various pathways, including residual cortico-spinal projections and the brainstem’s reticular formation for ICMS, as well as through the monosynaptic and polysynaptic proprioceptive circuits for EES ^23,29,39,61^.

The addition of sacral stimulation at the onset of the extension phase of the swing enhanced extension velocity without impacting step height. The observed increase in both extensor and flexor muscle activity during the stance and swing phase, respectively, suggests that S1 EES may influence the locomotor pattern by recruiting and facilitating lumbar circuits ^38,62^. Further experiments are needed to investigate the impact of S1 EES timing on leg movement modulation.

### Limitations of the study and clinical perspectives

This study highlights the potential for activating locomotor networks through distinctive spatial and temporal stimulation configurations involving multiple spinal and supraspinal sources. Furthermore, it underscores the importance of the coordinated interplay between cortical and sensory inputs in enhancing motor output robustness. While the precise mechanisms behind the effects of various EES configurations were not extensively examined in this study, the results show that substantial dragging reduction can be achieved with EES alone and in combination with ICMS. A limitation of our study is the use of 40-Hz L2 stimulation for long-term rehabilitation experiments, which may be suboptimal for long-term recovery. While we show that high-frequency stimulation is more effective for immediate movement modulation, optimizing the timing of spinal and cortical stimulation for plasticity remains critical. Our approach serves as a proof of concept, not claiming optimal timing, but highlighting the need for future research. As shown by several research groups ^43,46,52,58,59,63^, the timing of these stimulations is key to promoting plasticity, and even a 10-20 ms difference in the delay between cortical and spinal stimuli with respect to the optimal one can have radically different effects on long-term potentiation ^46^. Future studies should explore how these plasticity mechanisms can be integrated with our method to enhance movement recovery through better-timed combined stimulation.

Future preclinical investigations can expand on these findings to probe mechanisms and further explore the potential of combined neurostimulation strategies, potentially leading to distinct and functionally meaningful outcomes through the temporal and spatial modulation of different circuits using various stimulation parameters. Combining multiple complementary stimulation modalities holds promise for treating various injury conditions, including incomplete SCI and subcortical stroke. Integrating brain stimulation to reinforce volitional commands could enhance the effectiveness of EES and vice versa. The success of such strategies will depend on several non-exclusive factors that require further investigation, including: i) reinforcing volitional efforts during motor rehabilitation training to prime the lumbosacral circuitries and promote the reorganization of the entire chain of motor commands ^64,65^; ii) delivering stimulations at the precise timing when supraspinal and spinal inputs interact at the spinal level, optimizing facilitation and fostering associative plasticity ^43,46,58,59^; iii) accounting for the phase-dependent nature of motor responses induced by each stimulation source ^7,8,27,30,38,66^.

The timing of interventions after SCI might also be critical, with most rodent studies starting neuroprosthetic training during the acute recovery stage ^11,27^, contrasting with humans, where it typically begins at least a year post-injury. The impact of this timing discrepancy on responses to neurostimulation strategies for modulating locomotor networks requires further investigation, as does the assessment of cortico-spinal intervention efficacy across different time windows.

While the efficacy of EES and ICMS hinges on their specificity, the invasive nature of surgical procedures can restrict their widespread application. However, epidural electrodes have been effectively utilized for EES delivery in humans ^5,6,8,67^, and Utah arrays for ICMS have been tested successfully ^68^. Balancing effectiveness with invasiveness remains a challenge; intracortical electrodes offer high specificity but are invasive, while non-invasive brain stimulation may be less effective for precise movement modulation despite its suitability for conditioning protocols ^56^. New probe designs, including intravascular electrodes, have the potential to mitigate these issues ^69^.

Another challenge that multipronged interventions may face is the complexity of optimizing large parameter spaces. When combining two sources of stimulation, in this case cortical and spinal, the number of stimulation parameters options grows exponentially. Machine learning algorithms, such as Gaussian-Process-based Bayesian Optimization, provide a new approach for autonomously optimizing neurostimulation ^70,71^. This advancement is expected to aid in the clinical adoption of these techniques. With ongoing technological progress, we foresee that similar multimodal neuroprosthetic interventions will notably enhance functional outcomes and support personalized treatment strategies.

## Data and materials availability

The data generated throughout this study are available in a tabular format as supplementary material. Simple MATLAB code was used to draw figures from these data, and it is available from the authors upon reasonable request.

## Acknowledgments

The authors would like to thank Hugo Delivet-Mongrain, Elena Massai, Véronique Chouinard, Uliana Kaschii for their participation in animal handling, data acquisition, and analysis; Marjolaine Homier, Stéphane Ménard, Raphaël. Santamaria, and the staff at the Division des Animalerie for supporting our animal care.

## Funding

Natural Sciences and Engineering Research Council of Canada RGPIN-2015-03860 (MM)

Ministère de l’Économie, de l’Innovation et de l’Énergie du Québec PSOv2d-54829 (MM, MB)

New Frontiers in Research Fund Exploration NFRFE-2022-00394 (MB)

Fonds de Recherche Québec Santé (MM, MB).

MB was supported by postdoctoral fellowships from the Institut de valorisation des données (IVADO), the TransMedTech Institute, the FRQS.

RD was supported by a scholarship from IVADO and a departmental scholarship in memory of Tomás A. Reader.

DB was supported by a scholarship from the Institute TransMedTech.

RGH was supported by a scholarship from the Natural Sciences and Engineering Research Council of Canada and AS by a scholarship from the faculty of Medicine at the University of Montreal.

## Author contributions

Conceptualization: MM, MB

Methodology: MM, MB

Investigation: MM, MB, RD, RGH, AS

Formal analyses: MM, MB, RD, RGH, DB

Visualization: MM, MB, RD, DB

Validation: MM, MB, RD, RGH, DB

Data curation: MM, MB

Software: MB

Supervision: MM

Funding acquisition: MM, MB

Resources: MM

Project Administration: MM

Writing—original draft: RD, MM

Writing—review & editing: MM, MB, DB

## STAR METHODS

### Study design

The study’s sample size was determined through a power analysis using Monte Carlo simulation. An n=6 was established to achieve an 80% chance of reporting statistically significant results for true effect sizes twice the inter-subject variability σ (β=0.2, α=0.05, one-sided Wilcoxon rank-sum test), while accounting for a 50% chance of a non-responder within the group. For chronic effects on function recovery, we estimated variability based on previous experience with a baseline cohort (S.D. of 15% ladder score, assumed normally distributed), assuming a similar variability for the intervention cohort. The study aimed to report a +25% increase in performance using neuroprosthetic training. The chronic study was designed to include the minimum cohort size with at least an 80% chance of capturing the expected effect size to a significance threshold of α=0.05, corresponding to n=6. Only rats scoring 0-20% of ladder success one week post-SCI (pre-training) and without substantial deficits on the right hindlimb (a potential sign of over-hemisection) were included in the training study. After the study’s conclusion, lesion sizes were evaluated.

### Animals

All procedures followed the guidelines of the Canadian Council on Animal Care and were approved by the Comité de déontologie de l’expérimentation sur les animaux (CDEA; animal ethics committee) at Université de Montréal. Twenty-nine adult female Long-Evans rats weighting 270-330g (Charles River Laboratories, line 006) were used. Rats were provided *ad libitum* access to food and water throughout the duration of the experiments. The individual participation of each rat in the experiments is elaborated in Table S1.

### Surgical procedures

All surgical interventions were carried out under isoflurane general anesthesia, and local anesthesia (Lidocaine 2%) was administered at the incision sites. Analgesic (buprenorphine) and antibiotic (Baytril) medications were administered for 3-4 days following surgeries.

A first surgery was conducted to implant electromyographic (EMG), cortical and spinal electrodes. Differential EMG wires were inserted into the left and right tibialis anterior (ankle flexor) and medial gastrocnemius (ankle extensor), the biceps femoris (knee flexor) and quadriceps femoris (knee extensor) and in the left and right gluteus maximus (hip extensor). In the same surgery, a craniotomy was performed (approximately 3 mm x 2 mm) and the dura was removed from the right hindlimb motor cortex. A 32-channel array consisting of four columns spaced by 0.375 mm and eight rows spaced by 0.250 mm, (Tucker-Davis Technology, USA) was stereotaxically inserted into cortical layer V (1.5mm electrode depth), with the top left site of the array positioned at coordinates 1.1 posterior and 1.3 lateral from bregma, in the same coordinates of our previous study^27^. To optimize EES and engage both the lumbar and sacral spinal circuits in promoting stepping, two stimulating EES electrodes were sutured to the dura, positioned between the left dorsal roots and the spinal midline over the L2 and S1 spinal segments. A common ground wire (∼1 cm uninsulated at the distal end) was inserted subcutaneously around the torso. This ES configuration targeting L2-S1 has demonstrated superior efficacy compared to single-site stimulation, successfully inducing hindlimb stepping after complete SCI in recent rodent studies ^7,10,14,72^. Brain arrays, spinal and EMG wires connectors were all fixed using dental acrylic and microscrews secured to the skull for fixation. It’s important to note that in the long-term recovery experiments (Fig. 4), the rats did not all receive the same implants. Rats undergoing treadmill training only did not receive either cortical or spinal implants, while rats undergoing spinal stimulation did not receive cortical implants. Following completion of baseline experiments, a second surgical procedure was conducted to induce an incomplete SCI ^35^. T9 vertebra was exposed, followed by a partial laminectomy to expose the dorsal spinal cord. Lidocaine 2% was applied to reduce spinal reflexes before incising in the dura. A lateral hemisection of the left spinal cord was performed with a micro-scalpel. Gelfoam (Pfizer) was placed over the cord, followed by suturing of the muscle layers and skin. Manual bladder expression was employed as needed during the first week post-injury to assist proper micturition.

### Behavioral assessments

Rats’ locomotor performance was assessed in a treadmill, open field, and ladder. After acclimatization to handling, they were trained to walk quadrupedally on a treadmill at 23 cm/s using food rewards. While not trained for ladder or open field, occasional exposure facilitated acclimatization.

#### Treadmill walking

Each trial comprised the analysis of 10 consecutive steps during quadrupedal walking on a treadmill at 23 cm/s. Kinematics were recorded at a rate of 119.2 Hz, utilizing six reflective markers on key anatomical points: iliac crest, trochanter (hip), condyle (knee), malleolus (ankle), fifth metatarsal (foot), and fourth toe tip (limb end point). DeepLabCut ^73^ was employed for initial data processing, followed by manual curation to rectify any misidentifications. By automatized gait analysis, several kinematics parameters were used to evaluate locomotor performance. Step height was calculated by subtracting the average vertical position of the foot during stance from its maximum vertical displacement during swing. Flexion velocity refers to the maximum vertical speed of the foot during hindlimb flexion occurring in the swing phase. Extension velocity was defined as the maximum vertical speed (cm/s) achieved by the foot during leg extension until foot contact. Dragging was calculated as the percentage of the swing phase duration during which the foot moved forward while maintaining dorsal contact with the treadmill belt. 15 additional kinematic variables were used to perform multivariate analysis of gait via Principal Component Analysis (PCA), with the full list of variables and factor loadings indicated immediately next to the PCA representation (Fig. 6A). Blinding was not applied to kinematic analysis, which is nonetheless a highly automatized process.

#### Horizontal ladder

In rats subjected to chronic training, skilled voluntary locomotor control was assessed during ladder crossing. Rats were recorded at a frame rate of 100 frames per second while crossing a 130 cm horizontal ladder with rungs regularly positioned 2 cm apart. In each session, the results of five complete continuous crossings were averaged per each rat. Each trial consisted of approximately 10 steps. Performance was evaluated in a blinded manner by two experimenters, who calculated the percentage of faults made by the left foot over the total number of consecutive steps.

#### Open field

To assess the global locomotor and postural abilities, rats underwent assessment in an open field using neurological scoring scale developed in our previous study ^35^. Rats were recorded while engaging in 4 minutes of spontaneous locomotion within a circular Plexiglas arena (diameter: 96 cm, wall height: 40 cm) featuring an anti-skid floor. A locomotor score ranging from 0 to 20 was assigned by an experimenter blinded to the rats’ groups, considering specific parameters: 1) Articular movement amplitude of hip, knee, and ankle (0 = absent, 1 = slight, 2 = normal); 2) Stationary and active weight support of the limb (0 = absent, 1 = present); 3) Digit position of hindlimb (0 = flexed, 1 = atonic, 2 = extended); 4) Paw placement at initial contact (0 = dorsal, 1 = internal/external rotation, 2 = parallel); 5) Paw orientation during lift (1 = internal/external rotation, 2 = parallel); 6) Movement during swing (1 = irregular, 2 = regular); 7) Coordination between the fore– and hindlimb (0 = absent, 1 = occasional, 2 = frequent, 3 = consistent); and 8) tail position (0 = down, 1 = up).

### Neurostimulation delivery

Our neurostimulation approach, previously detailed in prior work ^27,30^, involves real-time processing of EMG activity during treadmill locomotion. Trigger events, identified by EMG signal crossing a manually set activation threshold, initiated cortical and/or spinal stimulation with specific delays. Before each experiment involving ICMS, the hindlimb motor cortex of rats was mapped to identify the stimulation channel eliciting clear hindlimb flexion at the lowest current threshold, for each experiment and each training session. ICMS consisted in 40 ms biphasic trains at 330 Hz (cathodic first, 200 μs/phase, 50-μs interphase interval, 20-300 μa). Spinal stimulation consisted in 40-250 ms biphasic trains at 20-330Hz (biphasic monopolar, 200 μs/phase, 50-μs interphase interval, 150-600 μa). The term “spatiotemporal” refers to the synchronization of neurostimulation delivery with a specific locomotor phase i.e. in phase-coherence, and at precise location, whether cortical, lumbar, and/or sacral. In this context, ICMS was delivered 40 ms before the lift onset, followed by L2 EES synchronized with the lift onset. Subsequently, 90 ms later, S1 EES was delivered during the second phase of the swing, corresponding to the onset of the first extension phase of the gait cycle. Stimulation intensities were linearly spaced within a functional range, defined from a minimum visible effect to a maximum comfortable value for the animal, determined by the animal’s willingness to perform the walking task with increasing stimulation to obtain food rewards. For each episode of locomotion, a single, fixed stimulation intensity was selected from within the defined range, and all intensity values within the range were systematically tested in subsequent episodes. In rats engaged in neuroprosthetic training (Fig. 4), stimulation intensities were set to 70-80% of the functional range to achieve optimal training effects while ensuring the rats’ comfort during the training sessions. In experiments focusing on the characterization of EES parameters, one parameter was randomly tested while the others were held constant. Detailed descriptions of the specific EES parameters can be found in the figure captions or the Results section.

### Histological assessments of lesion size

At the end of the experiments, rats were euthanized with pentobarbital and transcardially perfused with a solution of 0.2% heparin in 0.1 M phosphate-buffered saline (PBS; pH 7.4), followed by a 4% paraformaldehyde (PFA) in 0.1 M PBS (pH 7.4). A spinal segment spanning from T6 toT10 was extracted and postfixed for 24 h in PFA. The tissue was cryoprotected in a solution of 30% sucrose until it sunks. 40 µm thick coronal sections were cut using a cryosat. Every third section was mounted onto slides and stained with cresyl violet (0.5%) to visualize cell bodies in the spinal gray matter, and luxol fast blue (0.1%) to visualize myelin in the spinal white matter. Lesion extent was assessed by quantifying the percentage of damaged tissue in the coronal plan as described previously ^34,36^ (Fig. S10).

### Statistical analysis

For each statistical test, we initially assessed the normality of each data population. In cases where a population was deemed trivially non-normally distributed (e.g., instances where low amounts of dragging saturate to zero), non-parametric tests were utilized. For all other cases, a one-sample Kolmogorov-Smirnov test was employed to evaluate whether an assumption of normality is rejected by the dataset.

Statistical analyses are detailed in the caption of each Figure. Paired statistical comparisons were conducted using either Student’s t-test or the nonparametric Wilcoxon signed-rank test when normal distribution assumptions could not be made for at least one of the populations.

For nonpaired population samples, either Student’s t-test or the nonparametric Wilcoxon rank-sum test was applied, depending on the normality assumptions. When considering several hypotheses, the Bonferroni method was employed to correct the significance threshold.

For comparisons among multiple groups (Fig. 4, Fig. S4) or when replicates were individual muscles (Fig. S9), we employed one-way or repeated measures ANOVAs or the nonparametric Kruskall-Wallis test. Post-hoc t-tests were used for pairwise comparisons, and the statistical significance was adjusted using a Bonferroni correction.

We applied a threshold for statistical significance at *P*=0.05. Statistical analyses were performed using SPSS Statistics 29 or MATLAB.

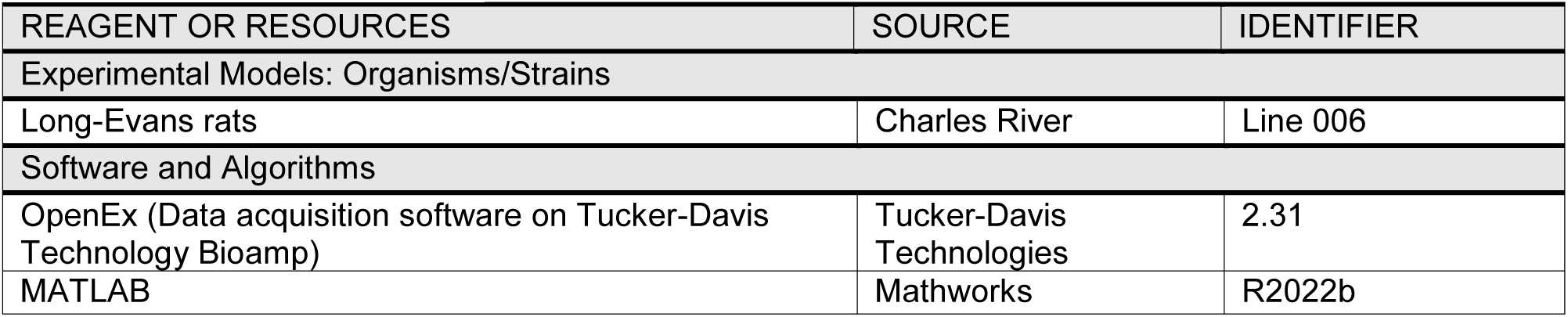
Key resources table.

## Supplementary information

### This PDF file includes

Figs. S1 to S10

Tables S1

Movies S1 and S2

**Fig. S1:**
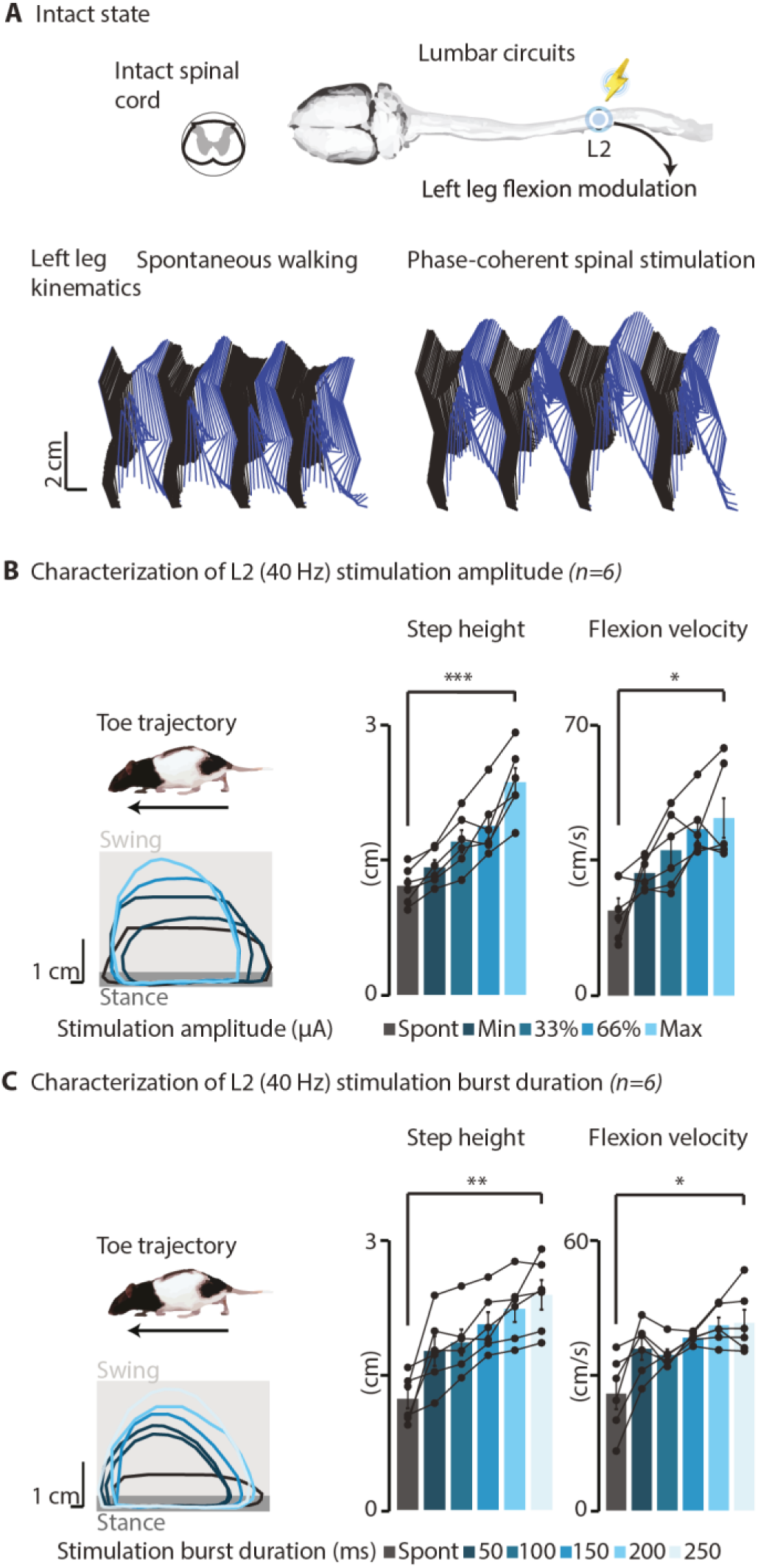
40Hz L2 EES intensity and duration modulate leg kinematics intact rats. (**A**), Schematic setup of EES delivery to the L2 spinal segment at the lift onset in intact rats. Stick diagrams depicting leg movements and toe trajectory without and with L2 EES in a representative rat. (**B**), Impact of EES intensity on leg kinematics. EES intensities were randomly tested within a functional range where the minimal intensity elicited a visible muscle twitch, and the maximal intensity was set to 90% of the maximal comfortable value for each rat (250-600 µA). EES burst duration was set at 120 ms. Step height and flexion velocity modulated with increased stimulation intensitys (step height: p= 0.0013, +95±9% of spontaneous walking, fit r^2^ min to max: 87±9%; flexion velocity: p=0.028, +132±48% of spontaneous walking, fit r^2^ min to max: 71±22%;). (**C**), Impact of EES burst duration (50-250ms range at maximal intensity captured in **B**) on leg kinematics. An increase in EES duration resulted in a linear increase in step height (p=0.003, +102±25% of spontaneous walking, fit r^2^ min to max: 90±3%). Increase in flexion velocity (p=0.015, +79±29% of spontaneous walking) was also observed. In all panels, individual data from n=6 rats are presented. Bar plots report mean values ± SEM. \**P*<0.05, \*\**P*<0.01, \*\*\**P*<0.001, paired, one-tailed t-test. Related to Fig. 2.

**Fig. S2:**
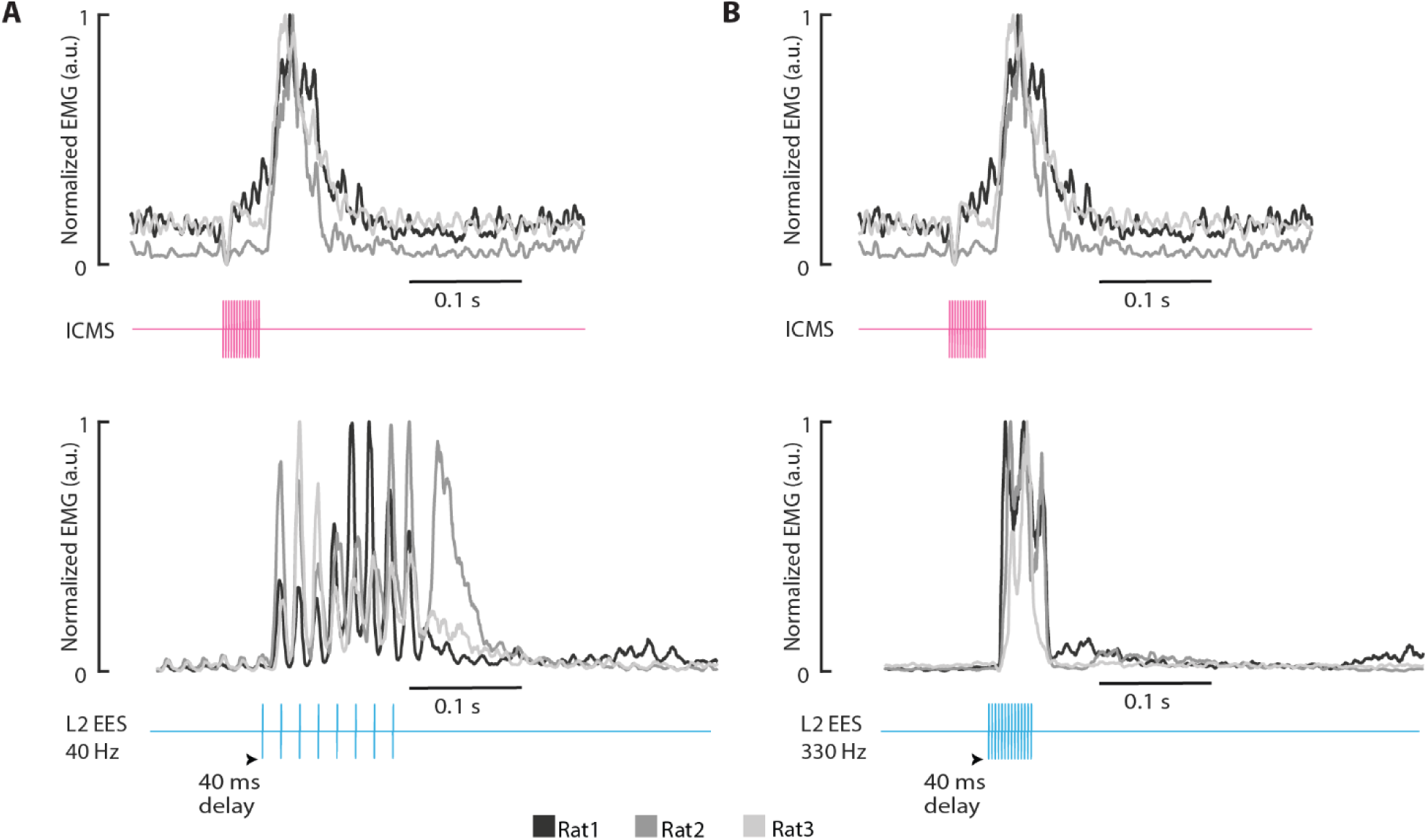
Impact of ICMS and L2 EES on ankle flexor motor evoked potentials. Motor evoked potentials recorded from the left ankle flexor (tibialis anterior) in response to ICMS and L2 EES at 40 Hz (**A**) and 330 Hz (**B**) in n=3 rats. The onset of L2 EES is delayed by 40 ms relative to ICMS, resulting in overlapping peak EMG volleys.

**Fig. S3:**
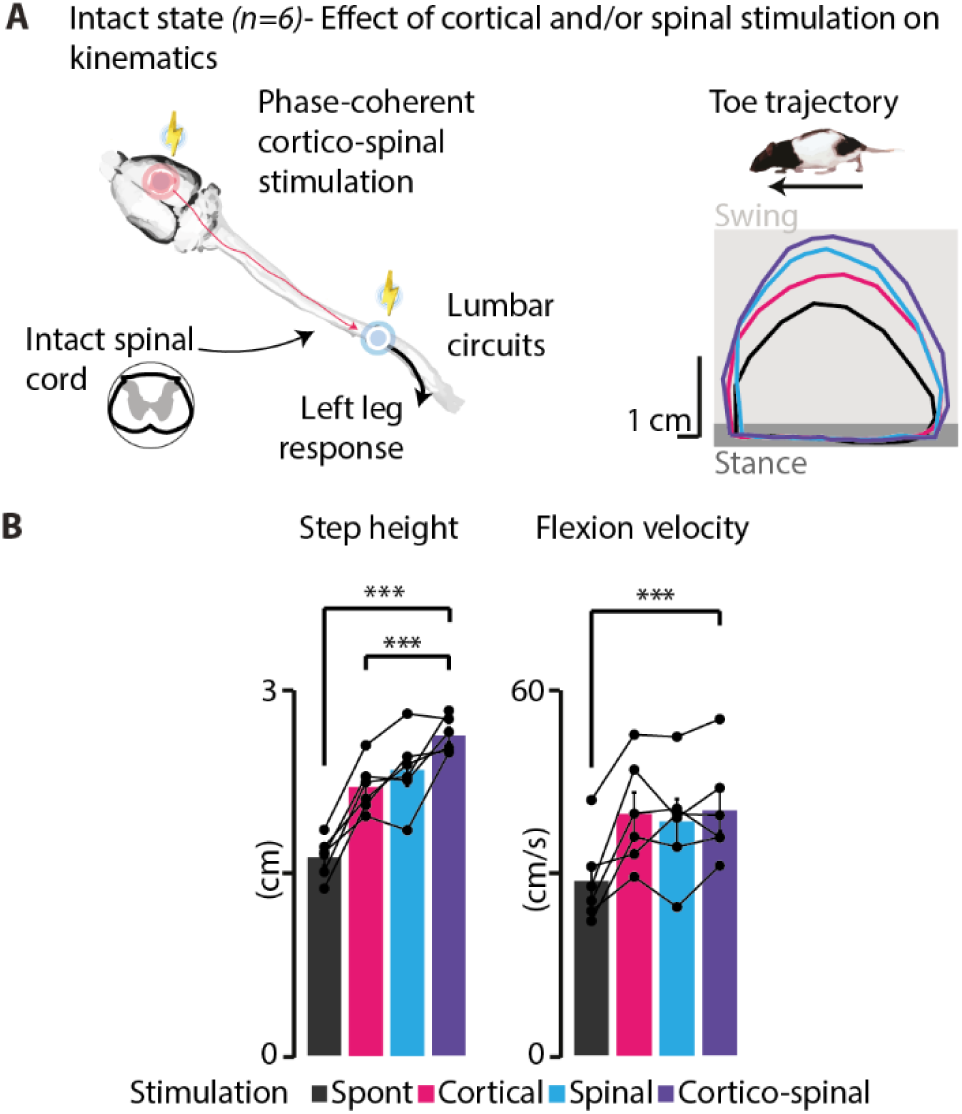
Combined cortico-spinal stimulation maximizes leg movements in intact rats. (**A**), Schematic setup of cortico-spinal stimulation delivery. Representative toe trajectory without and with cortical, spinal, and cortico-spinal stimulation., (**B**), Impact of individual and combined cortico-spinal stimulation on leg kinematics after SCI. Bar plots report mean values ± SEM (n=6 rats) of step height and flexion velocity for each condition of stimulation. Cortico-spinal stimulation significantly increased step height by 63±7% (p=0.0001) and flexion velocity by 42±6% (p=0.0006) compared to spontaneous walking. Cortico-spinal stimulation increased step height by +20±3% compared to independent cortical stimulation (p=0.0007). \**P*<0.05, \*\**P*<0.01, \*\*\**P*<0.001, paired, one-tailed t-test. The statistical significance threshold was adjusted for multiple comparisons. Related to Fig. 3.

**Fig. S4:**
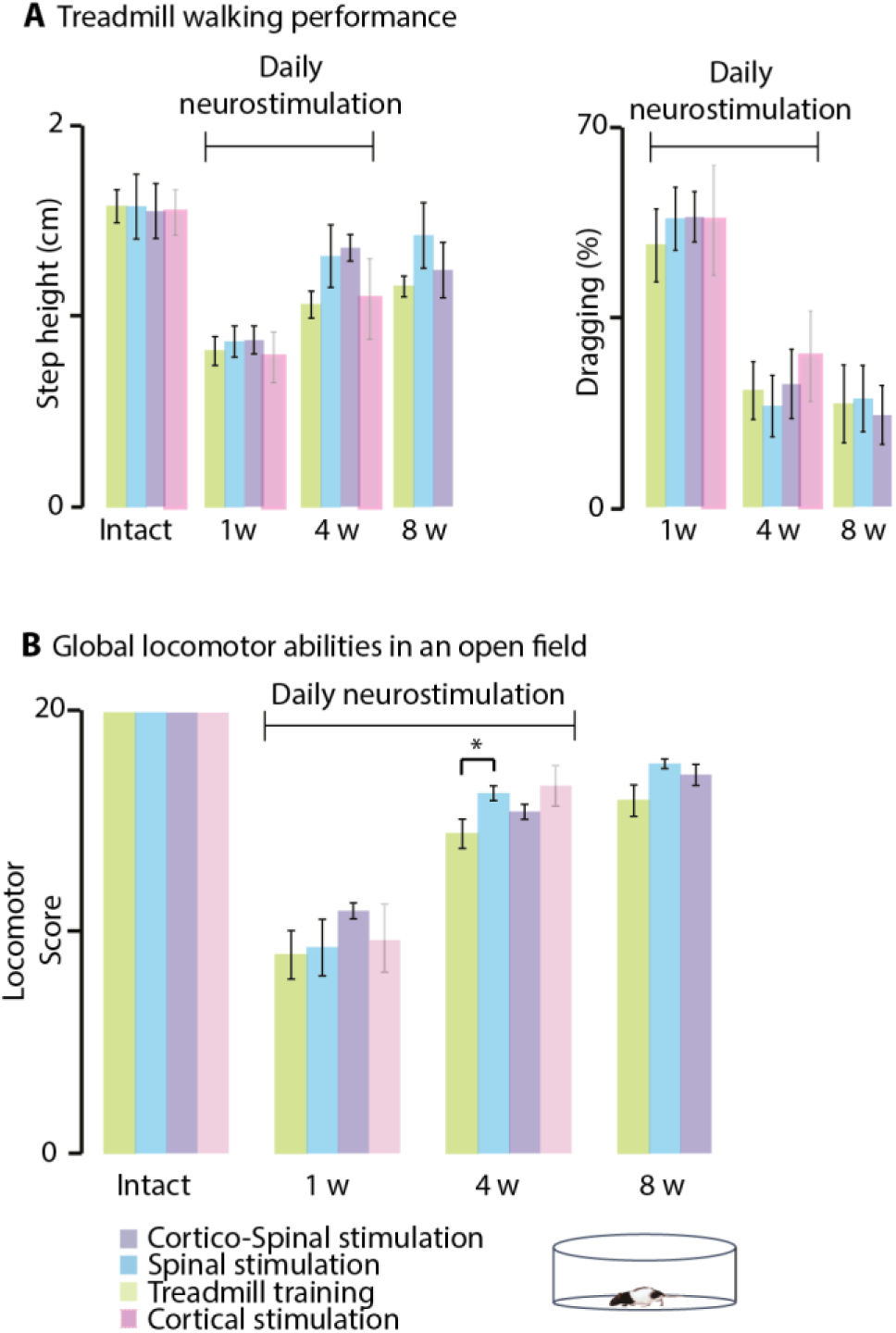
Treadmill performance and global locomotor abilities in rats submitted to locomotor training with or without cortical and/or spinal stimulation. (**A-B**), **Plots illustrate mean values** ± SEM of treadmill walking performance (**A**) and open field score (**B**) for each group (n=6 rats per group). (**A**), Step height and percentage of dragging were comparable across groups throughout the experiments (\**P*≥0.05, one-way ANOVA). (**B**), Global locomotor abilities were comparable across groups throughout the experiments expect at week 4, where the spinal stimulation group exhibited a higher score compared to the treadmill training group (*p=0.03, Kruskall-Wallis test supplemented with paired group comparisons). Data from rats subjected to cortical stimulation in our previous study ^27^ were included in the plots but were excluded from the statistical comparisons. Related to Fig. 4.

**Fig. S5:**
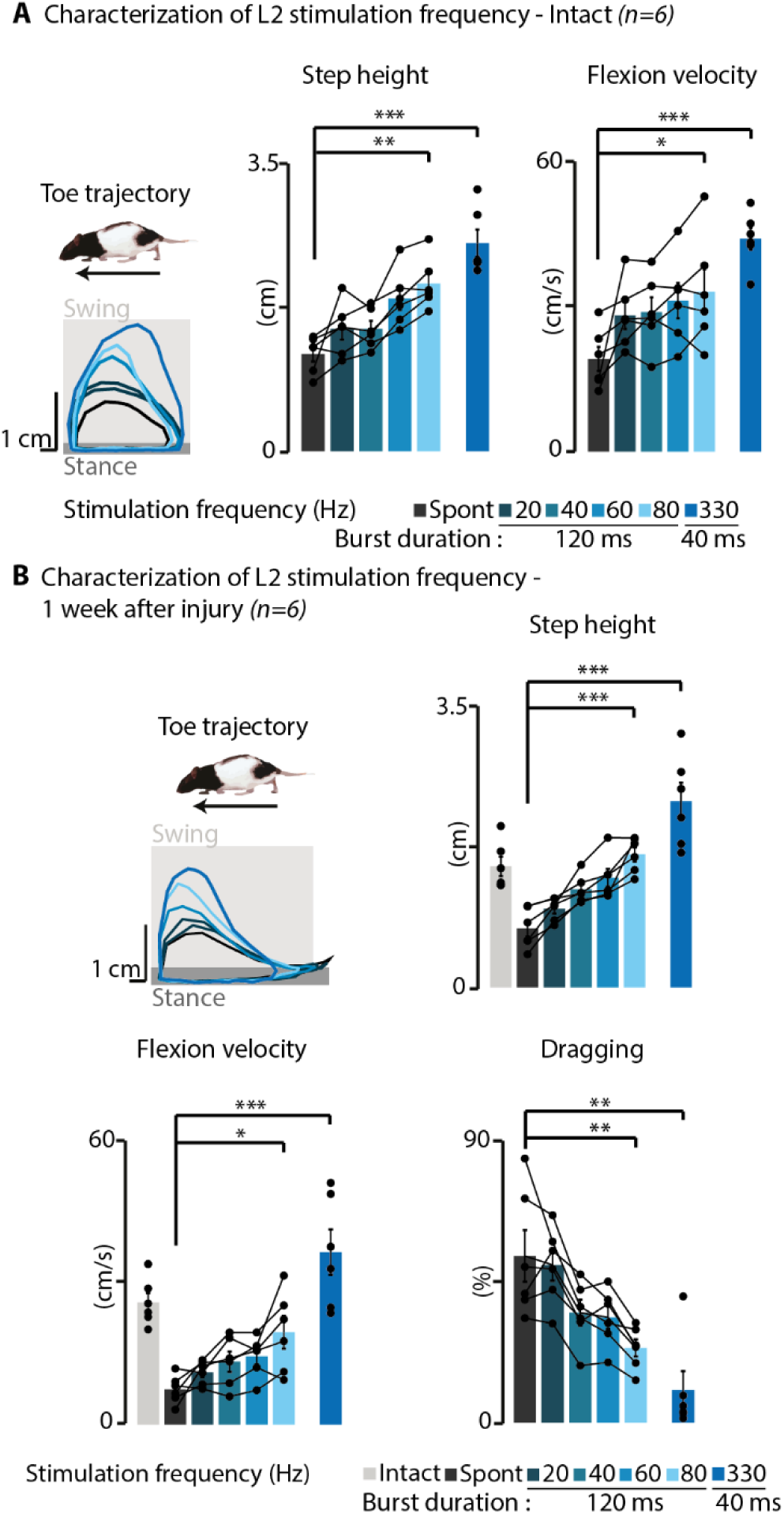
Impact of L2 EES frequency on leg kinematics in intact and SCI rats. (**A-B**), EES frequencies were randomly tested within a 20-80 Hz range with a fixed 120 ms duration and at 330 Hz with a fixed 40 ms duration. The maximal comfortable intensity for each rat before (**A**) and after SCI (**B**) was used. (**A**), In intact rats, increments in 120 ms EES frequency between 20-80 Hz resulted in a linear modulation of kinematic features, such as step height (fit r^2^ 20-80 Hz: 77±18%), with the highest modulation achieved at 80 Hz (p=0.007, +80±22% of spontaneous walking). The highest modulation occurred when the burst was reduced to 40 ms and the frequency increased to 330 Hz (spont vs 330Hz: step height: p=0.00003, +120±19%; flexion velocity: p=0.00003, +150±33%). (**B**), Similar results were observed after SCI, where EES frequency led to a linear increase in step height (fit r^2^ 20-80 Hz: 92±4%), with the highest trajectory achieved at 80 Hz (p=0.0006, +147±33% of spontaneous walking). 40 ms 330 Hz EES produced the maximal increase in step height (p=0.00009, +260±77% of spontaneous walking) and flexion velocity (p=0.00015, +534±203% of spontaneous walking), along with maximal decrease in dragging (p=0.009, –82±8% of spontaneous walking). Bar plots report mean values ± SEM (n=6 rats). \**P*<0.05, \*\**P*<0.01, \*\*\**P*<0.001, paired, one-tailed t-tests or Wilcoxon signed-rank tests. The statistical significance threshold was adjusted for multiple comparisons. Spont: spontaneous (no stimulation). Related to Fig. 5.

**Fig. S6:**
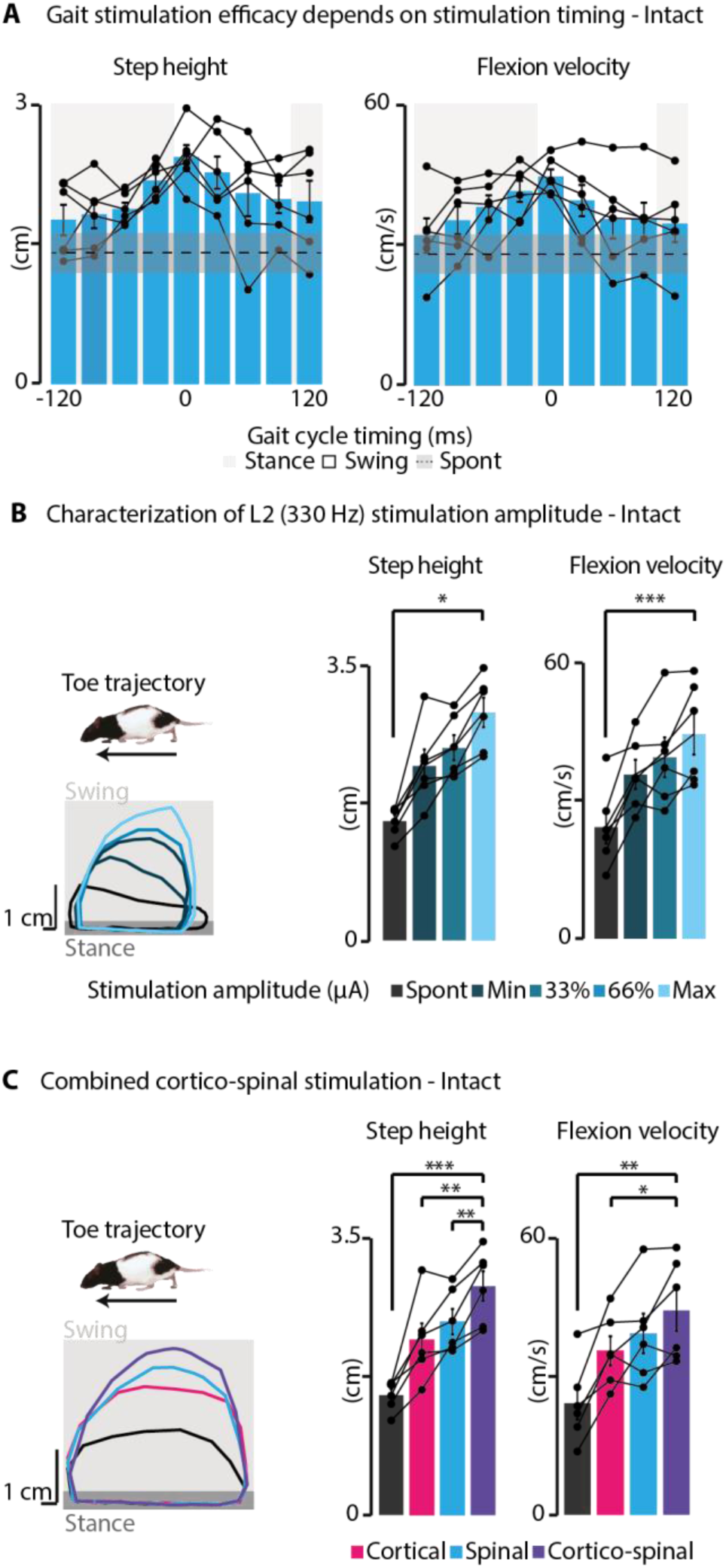
Characterization of cortico-spinal stimulation capabilities with high-frequency L2 EES on leg kinematics in intact rats. (**A**), Impact of high-frequency L2 EES (40 ms, 330Hz) timing on leg kinematics. Step height and flexion velocity are depicted for stimulation timings ranging from 120ms before to 120 ms after the lift, with 0 representing the lift onset. EES was most effective in modulating step height when delivered during the swing phase (stance vs swing: p=0.048, spont vs stance: p=0.002, spont vs swing: p= 0.002). (**B**), When delivered in synchrony with the lift, incremental increases in EES intensity (250 to 600 μA) proportionally modulated toe trajectory, leading to a linear increase in both step height (p=0.028, +124±26% of spontaneous, fit r^2^ min to max: 84±11%) and flexion velocity (p=0.0005, +131±31% of spontaneous walking, fit r^2^ min to max: 76±18%). (**C**), Combined cortical and high-frequency EES delivered at the lift onset maximized toe trajectory, significantly increasing step height by +91±8% (p=0.0003) and flexion velocity by +94±21 (p=0.005) compared to spontaneous walking. Compared to independent cortical and spinal stimulation, cortico-spinal stimulation increased step height by +33±7% (p=0.003) and 19±4% (p=0.007), respectively, and improved flexion velocity by 25±8% compared to cortical stimulation (p=0.04). Bar plots report mean values ± SEM (n=6 rats). \**P*<0.05, \*\**P*<0.01, \*\*\**P*<0.001, paired, two-tailed (A) and one-tailed (B-C) t-tests. The statistical significance threshold was adjusted for multiple comparisons. Spont: spontaneous (no stimulation). Related to Fig. 5.

**Fig. S7:**
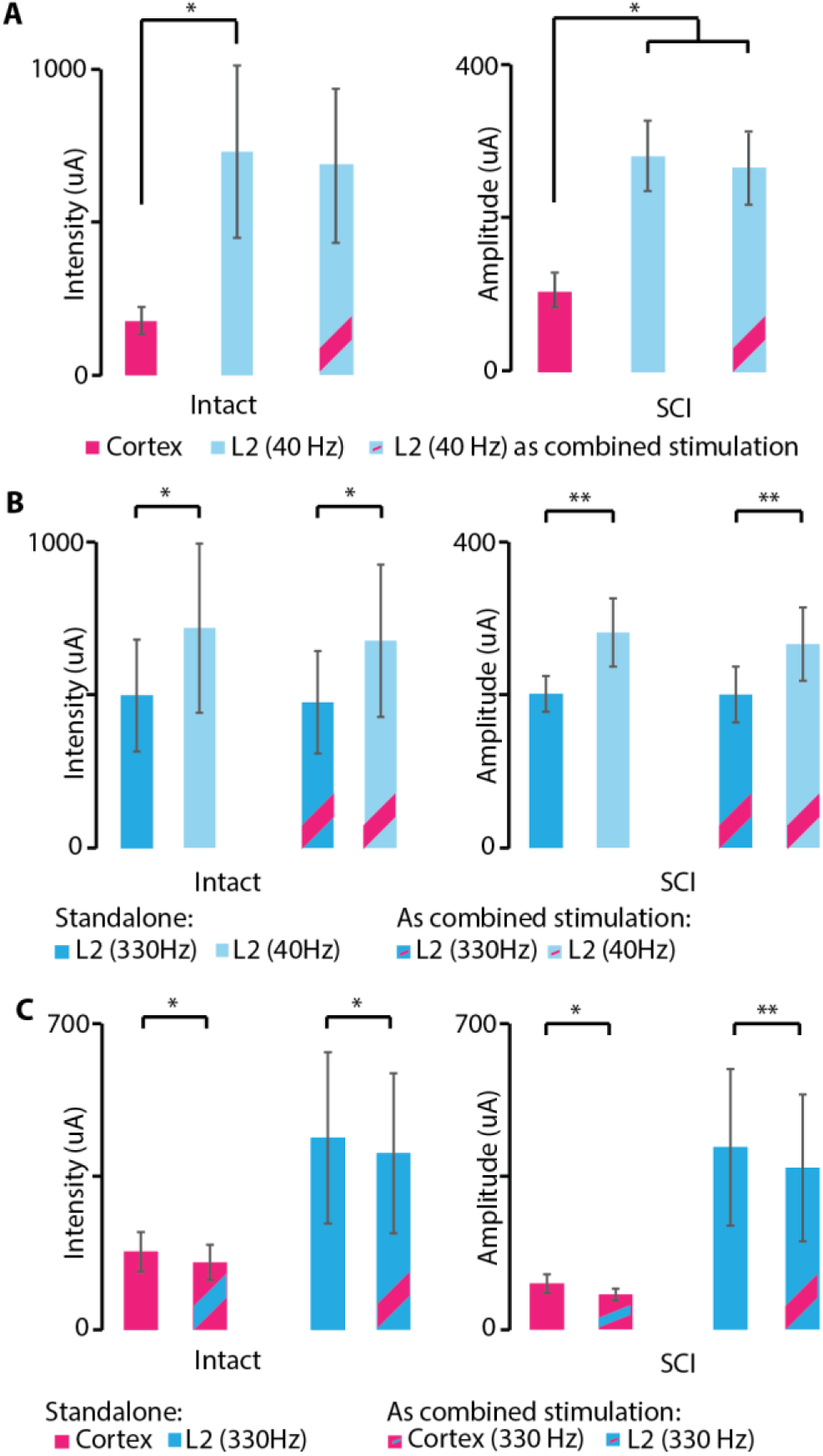
Maximal stimulation intensities used in intact and SCI rats. (**A**), Maximal comfortable intensities for individual and combined ICMS and L2 EES at 40 Hz. Cortical stimulation intensities were significantly lower than L2 EES at 40 Hz in both intact (–66 ± 12%, p=0.012) and SCI rats (–57± 15%, p=0.021), as well as when combined with L2 EES at 40 Hz in SCI rats (–55 ± 14%, p=0.037). (**B**), Maximal comfortable intensities used for standalone L2 EES stimulation at 330 Hz and 40 Hz frequencies, and when combined with ICMS. The intensities were significantly reduced by 25 ± 6% and 27 ± 4% when using L2 EES at 330 Hz compared to 40 Hz in intact (p=0.0443) and SCI rats (p=0.0090). The maximal intensities for L2 EES at 330 Hz when used as combined stimulation were also significantly lower compared to the 40-Hz EES used as combined stimulation, both in intact (–23 ± 6%, p=0.0439) and SCI rats (–24 ± 3%, p=0.0057). (**C**), Maximal intensities used for standalone ICMS and L2 EES at 330 Hz compared to the maximal intensities when combined. Intensities were significantly reduced by 13 ± 4% and 21 ± 5% when ICMS was paired with L2 EES at 330 Hz, compared to ICMS standalone in intact (p=0.0223) and SCI rats (p=0.0159). Similarly, intensities were significantly reduced by 9 ± 4% and 14 ± 2% when L2 EES at 330 Hz was combined with ICMS, compared to L2 EES alone in intact (p=0.0349) and SCI rats (p=0.0059). Bar plots report mean values ± SEM (n=6 rats). Different sets of n=6 rats were used for each panel. *P<0.05, **P<0.01, Friedman’s test (panel A, left), repeated measures ANOVA (panel B, right), paired, two-tailed t-tests (panels B-C). Related to Fig. 3, Fig. S3, Fig. 5 and Fig. S6.

**Fig. S8:**
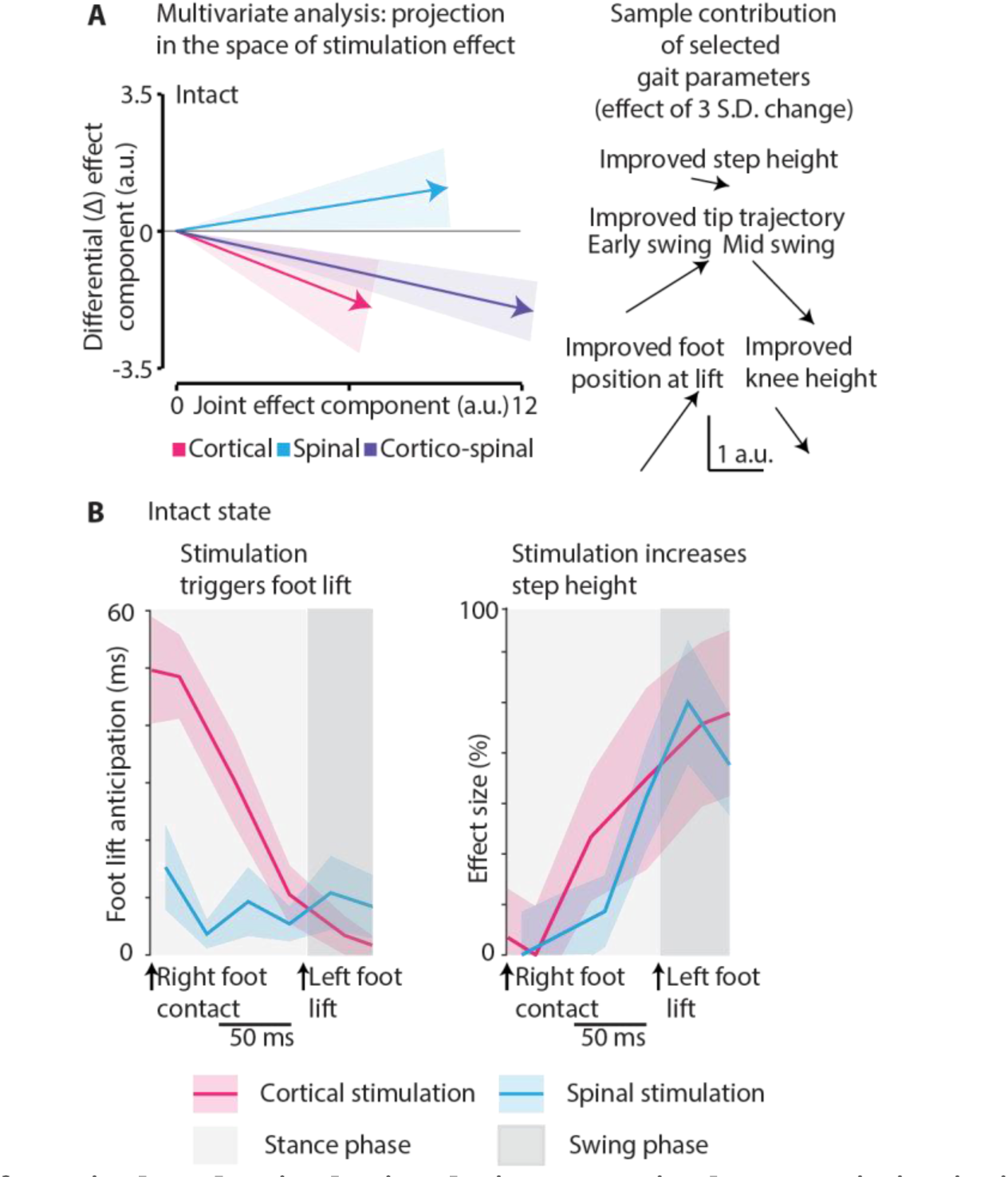
Impact of cortical and spinal stimulation on gait characteristics in intact rats. (**A**), Multivariate kinematic analysis qualitatively displays that combined cortico-spinal stimulation occupies an intermediate angular position between individual vectors, indicating a joint, and thus greater, overall impact on gait characteristics. (**B**), While both L2 EES and cortical stimulation increase step height, only ICMS initiates the swing phase. Shaded areas in graphs represent inter-individual variability. Related to Fig. 6.

**Fig. S9.**
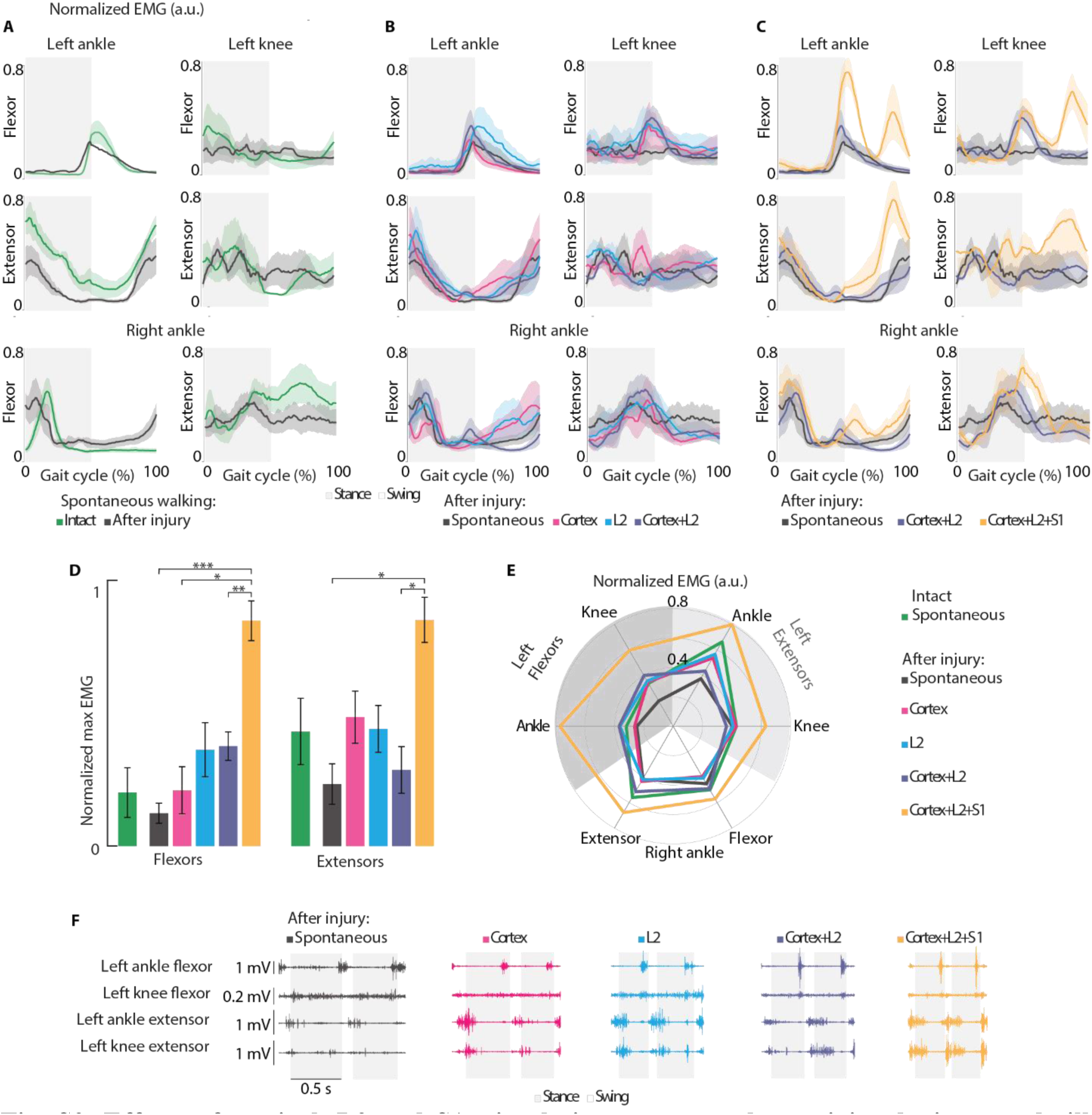
Effects of cortical, L2 and S1 stimulations on muscular activity during treadmill walking. (**A-C**), Average normalized EMG activity pattern for six recorded muscles: ankle and knee flexors and extensors in the left leg, and right ankle flexor and extensor. (**A**), Comparison between intact and SCI rats. (**B**), In SCI rats, comparison between spontaneous walking, individual and combined cortical and spinal stimulation. (**C**), In SCI rats, visualization of the EMG response to added S1 stimulation. (**D**), Bar diagram indicating peak normalized EMG activity for flexor and extensor muscles across conditions. (**E**), Spider plot of peak normalized EMG activity for individual muscles. (**F**), Sample muscle traces across stimulation conditions for one rat. \**P*<0.05, \*\**P*<0.01, \*\*\**P*<0.001, repeated measures ANOVA supplemented by Tukey’s HSD post-hoc test. The statistical significance threshold was adjusted for multiple comparisons. Spont: spontaneous (no stimulation). Related to Fig. 7.

**Fig. S10:**
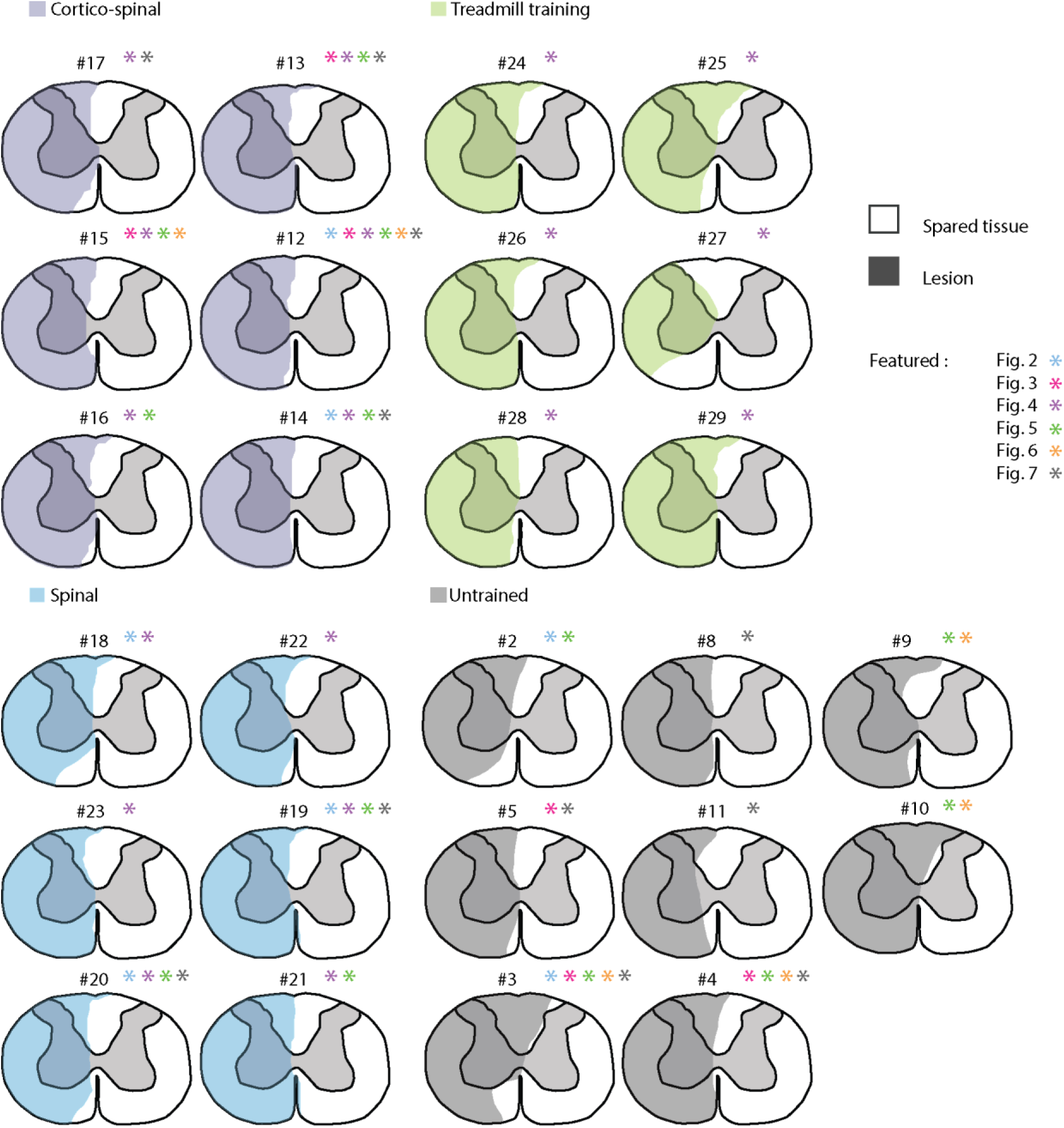
Spinal lesion severity. Hemisection profiles at the epicenter level for each rat included in the study. Related to Fig. 2-7.

**Table S1:**
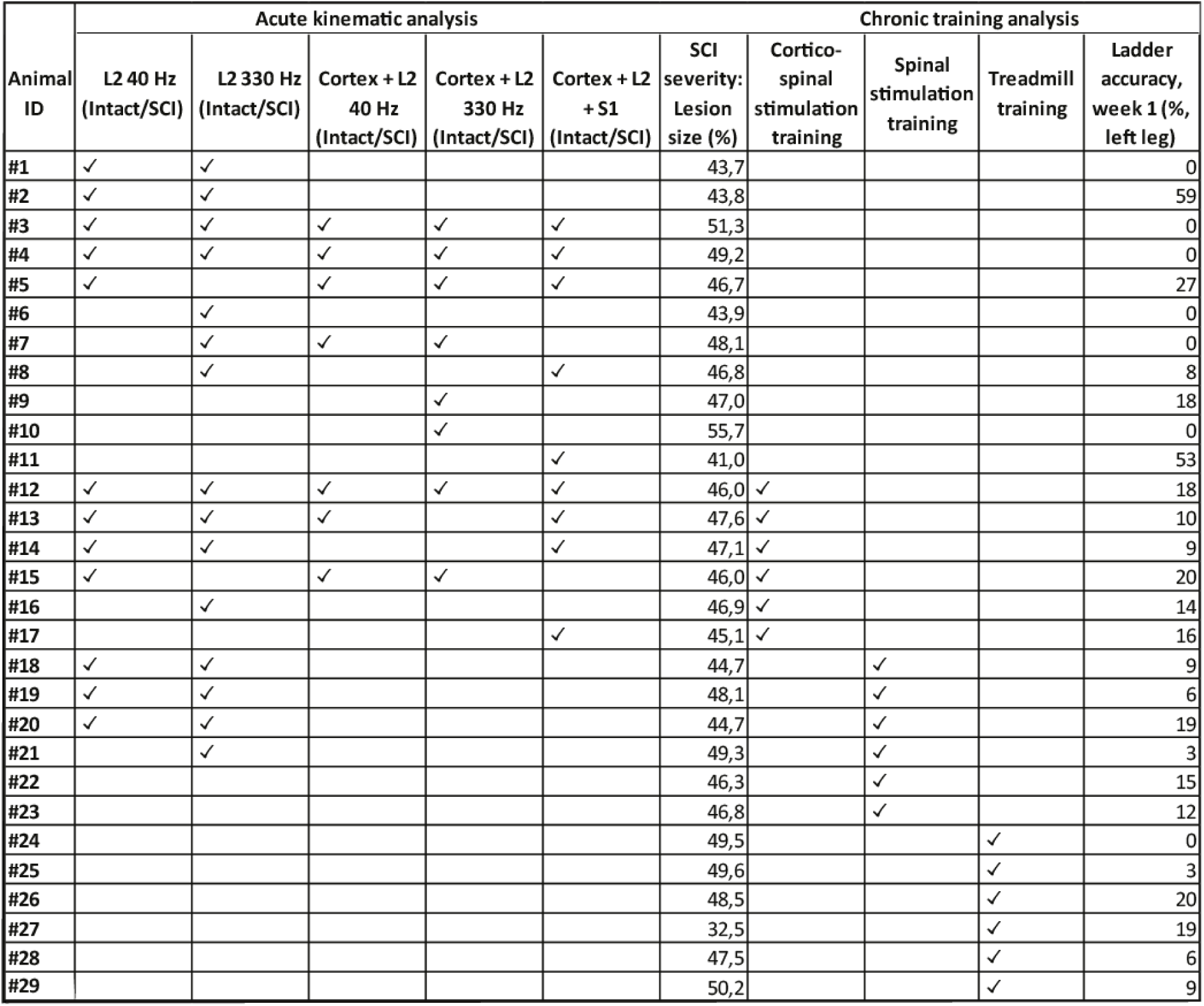
Individual data of rats included in the study. A total of 29 rats were involved in the study. Twenty-one rats participated in the acute experiments to test the immediate effects of stimulation protocols. Some of these rats were also used in the chronic experiments to assess the long-term effects of L2 EES or combined ICMS and L2 EES. The chronic experiments included 6 rats per group, for a total of 18 rats.

**<u>Movie S1</u>.**

Treadmill locomotion during spontaneous walking, with L2 EES, cortical stimulation, and combined cortico-spinal stimulation in the same rat nine days after SCI. EES was delivered at 40 Hz and cortical stimulation at 330 Hz. Playback speed has been slowed down 4x. Related to Fig. 5.

**<u>Movie S2</u>.**

Treadmill locomotion during spontaneous walking, incorporating L2 EES, combined cortical and L2 EES stimulation, and combined cortical, L2, and S1 EES in the same rat nine days after SCI. All stimulations were delivered at 330 Hz. Playback speed has been slowed down 4x. Related to Fig. 7.

